# Mapping the developmental potential of mouse inner ear organoids at single-cell resolution

**DOI:** 10.1101/2023.09.08.556845

**Authors:** Joerg Waldhaus, Linghua Jiang, Liqian Liu, Jie Liu, Robert Keith Duncan

**Author notes:** Senior Author (Joerg Waldhaus, Jie Liu, Robert Keith Duncan). Authors contributed equally (Joerg Waldhaus, Linghua Jiang). Corresponding author, Medical Sciences I Building, Rm. 5315C, 1150 West Medical Center Drive, Ann Arbor, MI 48109-5616, USA.

## Abstract

Inner ear organoids recapitulate development and are intended to generate cell types of the otic lineage for applications such as basic science research and cell replacement strategies. Here, we use single-cell sequencing to study the cellular heterogeneity of late-stage mouse inner ear organoid sensory epithelia, which we validated by comparison with data sets of the mouse cochlea and vestibular epithelia. We resolved supporting cell sub-types, cochlear like hair cells, and vestibular Type I and Type II like hair cells. While cochlear like hair cells aligned best with an outer hair cell trajectory, vestibular like hair cells followed developmental trajectories similar to *in vivo* programs branching into Type II and then Type I extrastriolar hair cells. These results highlight the transcriptional accuracy of the organoid developmental program but will also inform future strategies to improve synaptic connectivity and regional specification.

## Introduction

The inner ear, located in the temporal bone of the skull, is divided into two anatomical compartments: the snail shell shaped cochlea that transduces sound and the vestibular apparatus detecting linear and rotational acceleration. Both organs develop from a shared anlage, the otocyst; and while the ventral side of the otocyst gives rise to the cochlea, the vestibulum originates from the dorsal half.^1^ Even though both organs have different functional roles, they both rely on mechanosensitive hair cells (HCs) that convert mechanical stimuli into graded receptor potentials. At the cellular and molecular level, vestibular and cochlear HCs share many characteristics, such as the role of the hair bundles in detecting the stimuli or the function of synaptic proteins in signal transmission.^2^ Furthermore, the sensory cells of both organs are embedded in a layer of supporting cells (SCs) that, among other things, provide structural integrity and metabolic support.

However, there are numerous differences in anatomy and function of the HCs that relate to the different organs. For example, two types of HCs are characteristic for the organ of Corti, where one row of inner HCs functions to detect sound and three rows of outer HCs have a role in signal amplification.^3^ In comparison, HCs of the vestibular apparatus segregate into Type I and Type II HCs.^4^ They are characterized by different types of innervations and expression of marker genes such as *Ocm*, *Anxa4*, and *Spp1.*^5, 6^ Within these four distinct categories of HCs, there arises an array of more subtle structural, molecular, and functional differences by region within each organ.^7–10^

Genetic mutations, drugs, and environmental factors such as noise exert a negative impact on inner ear function.^11–13^ Damage that results in HC loss is irreversible and contributes to the prevalence of hearing and balance disorders worldwide.^14, 15^ In recent decades, advances in stem cell biology have opened new avenues to therapeutic discovery and novel approaches to regenerative medicine. One of the most significant advances in this area is the development of organoid culture platforms. Organoids are three-dimensional, multicellular model tissues that recapitulate many of the developmental, structural, and functional features of complex organs *in vitro.*^16^ As such, organoid platforms provide unique opportunities to explore the molecular mechanisms of disease and to generate cells or tissues for regenerative medicine.^17^ Koehler et al. established a systematic approach for the derivation of inner ear organoids,^18, 19^ where application of an artificial matrix (Matrigel) and manipulation of BMP, TGFβ, FGF, and WNT pathways result in the differentiation of sensory epithelia containing HC like cells (HCLCs), SC like cells (SCLCs), and synaptic connections to otocyst-derived neurons.^20^ More recently, inner ear organoid cultures have been used to model the genetic basis of inner ear disorders^21, 22^ and the developmental signals that regulate regional specializations in the ear.^23, 24^ However, the degree to which inner ear organoids mimic the complexity of the normal inner ear remains uncertain, limiting their broad utility in downstream applications.

Here we aimed to characterize the cellular diversity of organoid derived HCs at a whole transcriptome scale in an unbiased approach. Therefore, we used single cell RNA sequencing (scRNA-seq) and sampled 15,001 cells from late-stage mouse organoids at three different time points. HCLC and SCLC clusters were found and comparison with *in vivo* differentiated HCs revealed differentiation of auditory and vestibular like HCLCs *in vitro*. Finally, trajectory-based analysis of the vestibular like fraction of cells confirmed that organoid derived HCLCs develop into Type I and Type II like vestibular HCLCs. Importantly, gene expression dynamics along the trajectory closely resembled temporal changes during *in vivo* development, not only for a limited number of genes, but also at whole transcriptome level.

## Results

### Generation of inner ear organoids from *Atoh1*-nGFP mouse ESCs

For generation of inner ear organoids from mouse ESCs, cells were exposed to a combination of growth factors and small molecules (Figure 1A).^19^ We used an *Atoh1*-nGFP^25^ ESC line to label nascent HCs and to allow for enrichment of HCLCs. Cystic organoids were produced by approximately 40-60% of the aggregates in any culture plate, with the majority of those cysts incorporating GFP-positive cells indicative of the presence of sensory hair cell lineages (Supplemental Figures S1A-B). The experimental paradigm included dissociation of whole organoids, fluorescence activated cell sorting (FACS), and subsequent scRNA-seq analysis (Figure 1B). To verify that sensory HCLCs survived the isolation procedures, we stained the single-cell suspension from organoids 22 days-in-vitro (DIV) with an antibody to GFP and labeled the actin-rich hair bundles using a fluorescently conjugated phalloidin (Figure 1C).

**Figure 1.**
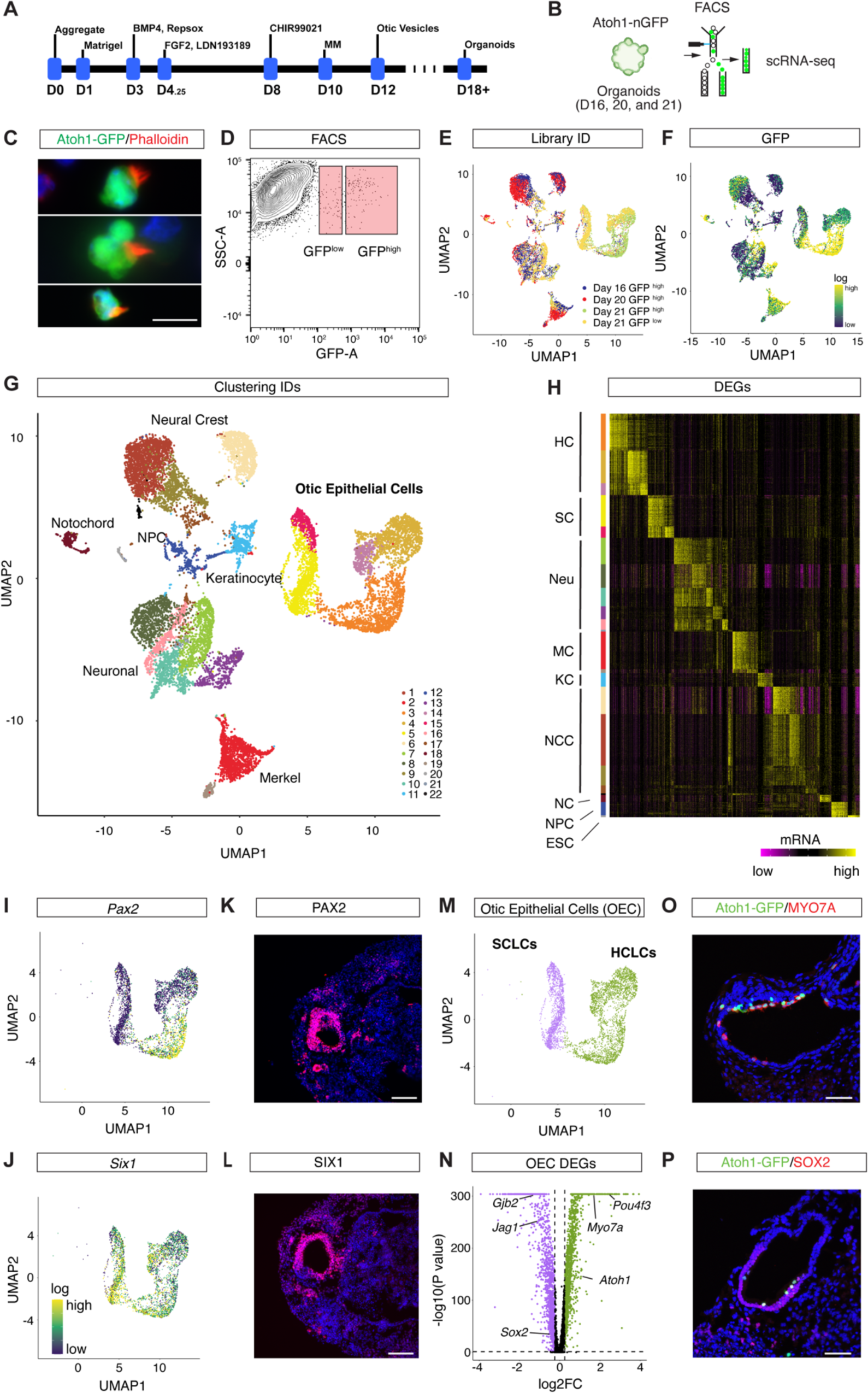
scRNA-seq profiling of late-stage inner ear organoids. (A) Differentiation strategy for inner ear organoids modified from Koehler et al. (B) Schematic representation of the experimental workflow used in this study. (C) Representative micrographs of individual HCLCs stained with an antibody against GFP and phalloidin in single cell preparations of D22 organoids. (D) FACS plot and gating strategy to isolate cells expressing GFP at high and low levels. (E) UMAP plots to show the clustering of 16, 20, and 21 DIV cells processed in four aggregated libraries to rule out technical variations for scRNA-seq experiments. (F) UMAP of all cells isolated with *Gfp* expression levels color coded. (G) UMAP plots of cells isolated with cell type annotations. Cell types constituting the otic epithelium segregated into 5 clusters. (H) Expression heat map for 15,001 organoid cells (y-axis) and DEGs (x-axis). Shown are the top 100 DEGs for each of the 11 clusters identified. Cluster identities were determined based on DEGs known as canonical markers (also see Supplemental Fig. S1E) (I,J) UMAP of all otic epithelial cells. *Pax2* (I) and *Six1* (J) mRNA expression projected. (K,L) Representative micrographs of 12 DIV cysts represented in cryosections and stained using antibodies. Anti-PAX2 (K) and Anti-SIX1 (L) counterstained with DAPI. (M) Otic epithelial cells segregated into 3,029 HCLCs (green) and 1,581 SCLCs (purple). (N) Volcano plots of DEGs comparing HCLCs and SCLCs. Cutoff: adjusted p-value < 0.05 and absolute value of log2FC > 0.25. (O,P) Representative micrograph of 22 DIV cysts represented in cryosections and stained using antibodies against GFP (O,P), MYO7A (O), and SOX2 (P). Scale bars: 10 µm in C, 100 µm in K,L, and 50 µm O,P. (B) was created with BioRender.com.

### Transcriptional profiling of inner ear organoids

Single cells from late stage whole organoid preparations at 16, 20, and 21 DIV were sorted based on GFP-fluorescence intensity levels into GFP-high and -low groups (Figure 1D and Supplemental Figure S1C) and processed individually for library preparation using the 10x Genomics scRNA-seq platform. Overall, 15,001 cells passed a stringent quality control (Figure 1E and Supplemental Figure S1D), and the sorting strategy was validated by plotting *Gfp* transcript levels onto the UMAP projection (Figure 1F). In total, 22 clusters (Figure 1G) were identified using the Seurat v3 ^26^ pipeline and differentially expressed genes (DEGs) (adjusted p-value < 0.05) were determined by comparing averaged gene expression for each cluster with the remaining cells (Figure 1H). Altogether, 23,356 DEGs were identified with an average count of 1,062 DEGs per cluster. Known marker genes among the DEGs were used to annotate cluster identities (Supplemental Figure S1E). Briefly, transcripts for early otic epithelial markers such as *Pax2* and *Six1* were detected in inner ear organoids between 16 and 21 DIV (Figures 1I,J). Similarly, antibodies raised against PAX2 and SIX1 stained the organoid vesicle intermediates at 12 DIV (Figures 1K,L). Furthermore, a separation of HCLCs and SCLCs (Figures 1M) was found based on differential expression of markers such as *Atoh1*, *Pou4f3*, *Myo7a*^27–29^ and *Sox2*, *Jag1*, and *Gjb2*^30–32^ (Figure 1N), respectively. Cryosections through the cysts reveal large, fluid-filled lumens lined with cells positively stained for the sensory HC marker MYO7A (Figure 1O). Among 10 independent organoids, over 90% of MYO7A-positive cells co-expressed GFP. Notably, organoids also contained SOX2-positive SCLCs, which occasionally co-labeled with GFP (Figure 1P). In summary, using scRNA-seq we successfully collected whole transcriptome profiles of 3,029 HCLCs and 1,581 SCLCs from three independent experiments across 5 clusters.

The sorting strategy resulted in collection of off-target cells that were excluded from further analysis but annotated based on the DEGs (Supplemental Figure S1E). Briefly, different types of neuronal like cells were represented in three clusters that expressed markers such as *Map2* and *Neurod2.*^33–35^ Merkel cell like cells were identified based on *Krt18*, *Syn2*, and *Piezo2*^36, 37^ expression, while a population of keratinocyte like cells was identified based on *Krt5*, *Krt14*, *Krt15*^38^ expression. Neural crest like cells were characterized by differential expression of *Twist1* and *Col1a2.*^39, 40^ Smaller clusters of notochord like cells, neural precursors, and residual pluripotent stem cells were identified by *T*, *Ascl1* and *Pou5f1* expression.^41–43^

### *In vitro* differentiation of SCLCs and non-sensory epithelial cells

We identified a large number of *Sox10*+^31^ (Figure 2A) SCLCs in our cell samples. This observation concurred with previous reports of low *Atoh1*-GFP expression levels in subpopulations of organ of Corti SCs such as inner border cells^44^ and is likely due to specifics of the transgenic reporter construct that utilizes a single *Atoh1* enhancer element to drive the GFP expression.^25^ The initial clustering resolved two sub-populations of SCLCs (Figure 2B). Based on differential gene expression (Figure 2C), sub-clusters were annotated as sensory SCLCs and non-sensory epithelial cells (NSECs).^45^ For example, the majority of SCLCs expressed markers such as *Sox2*, *Jag1*, and *Lfng* (Figures 2C,D and Supplemental Figure S1E),^32, 46, 47^ reminiscent of SCs in the sensory epithelia of the inner ear, whereas cells of the smaller cluster expressed genes such as *Oc90*, *Otol1* and *Otoa* (Figure 2C,E and Supplemental Figure S1E),^48–50^ resembling the cell types lining the inner ear fluid spaces outside the sensory epithelia. During embryonic development of the cochlea, these cell types constitute the cochlear roof, which will later give rise to structures such as stria vascularis and Reissner’s membrane. In summary, this finding demonstrates that organoid derived SCLCs differentiate into at least two different types of inner ear SCLCs, raising questions about the cellular identities of the different HCLC sub-clusters identified.

**Figure 2.**
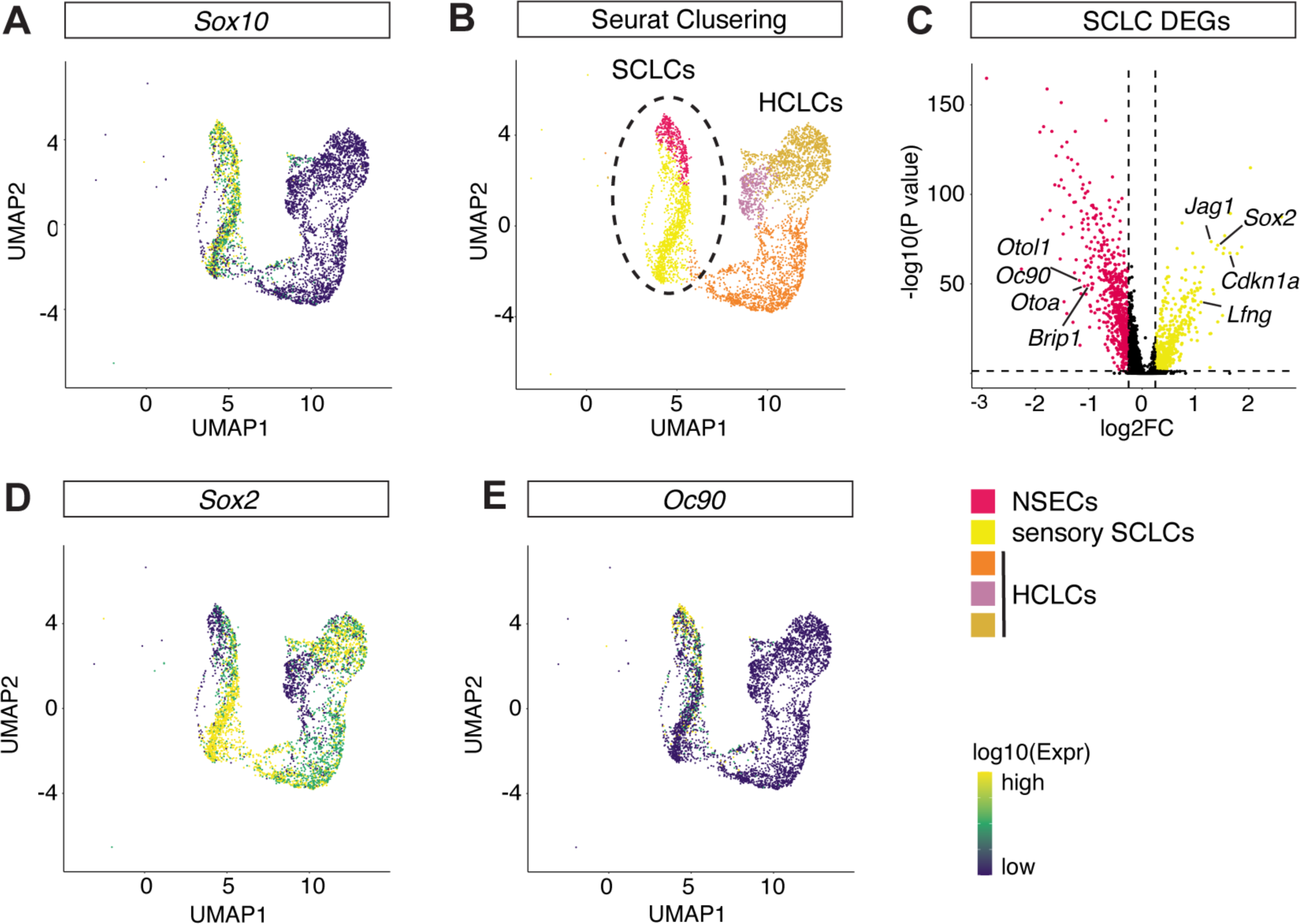
SCLC sub-cluster analysis. (A) UMAP of all otic epithelial cells with *Sox10* mRNA expression projected. (B) Seurat clustering resolved two sub-cluster in the SCLC population. (C) Volcano plots of DEGs comparing NSECs (red) and sensory SCLCs (yellow). Cutoff: adjusted p-value < 0.05 and absolute value of log2FC > 0.25. (D,E) UMAP of all otic epithelial cells with *Sox2* (D) and *Oc90* (E) mRNA expression projected.

### Transcriptional kinetics during *in vitro* HC differentiation

While SCLCs segregated based on spatial identity, an RNA velocity analysis^51^ suggests that the HCLC clustering corresponds to different states of HC maturation (Figure 3A). Briefly, the algorithm determines RNA velocity as a ratio of preprocessed versus processed mRNAs and allows for embedded projection of RNA velocity with UMAP clustering. Dynamically changing cell states are characterized by high RNA velocity, while differentiated states exhibit steady state kinetics with low RNA velocity. Analyzing transcriptional kinetics of *in vitro* generated HCs, we found differential RNA velocity within the three HC clusters (Figure 3A). For example, the cluster exhibiting highest RNA velocity was characterized by genes indicating early HC development such as *Atoh1*, *Lhx3*, and *Jag2* (Figures 3A-D).^29, 52, 53^ Whereas lower RNA velocities were associated with clusters that differentially expressed genes contributing to function of HCs such as *Tmc1*, *Espn* and *Kcna10* (Figures 3A-C,E).^54–56^ In summary, the RNA velocity analysis supports the hypothesis that HCLCs clustering along UMAP2 represent the temporal aspect of HC maturation as visualized in a pseudotime projection (Figure 3F).

**Figure 3.**
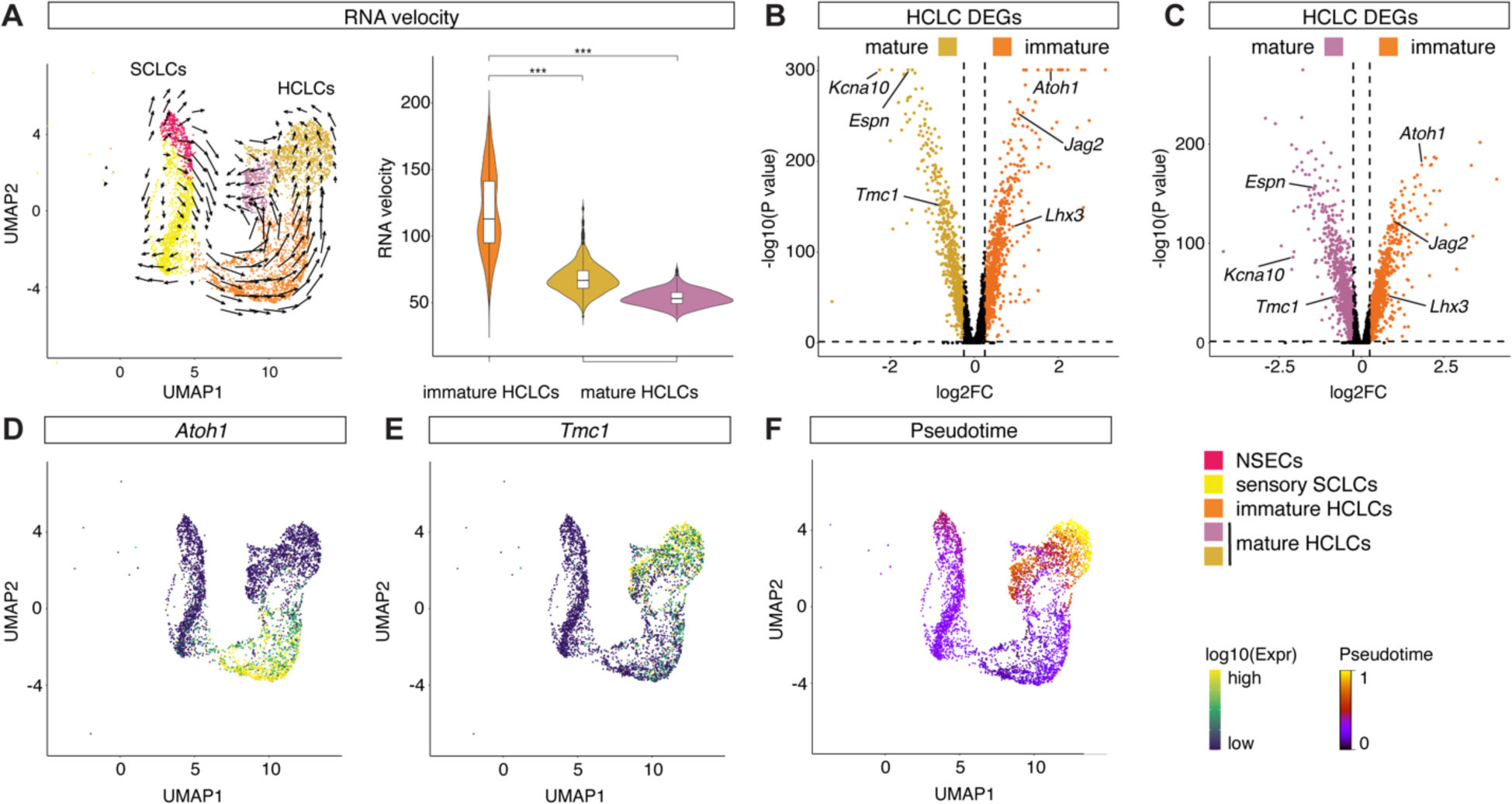
Transcriptional kinetics in HCLCs. (A) UMAP of all otic epithelial cells with cluster ID and RNA velocity projected as arrows (left panel). Length of the arrow corresponds to RNA velocity. Violin plots of RNA velocity for the three HCLC clusters (right panel). Cluster ID is color coded. The violin plots show significantly different RNA velocities between the immature cluster and the two mature clusters. ***P < 0.001 (Wilcoxon rank sum test). (B,C) Volcano plots of DEGs comparing immature (orange) and mature HCLCs (brown (B) and purple (C)). Cutoff: adjusted p-value < 0.05 and absolute value of log2FC > 0.25. (D,E) UMAP of all otic epithelial cells with *Atoh1* (D) and *Tmc1* (E) mRNA expression projected. (F) UMAP of all otic epithelial cells with RNA velocity based pseudotime superimposed.

### Differentiation of vestibular and auditory HCLCs and SCLCs *in vitro*

During inner ear organogenesis, auditory and vestibular HC lineages develop from opposite ends of the otocyst. Aiming to identify the lineage identities of the three HCLC clusters, we first re-clustered the HCLCs and SCLCs to increase cluster resolution using CellTrails^57^ and 3 SCLC and 5 HCLC CellTrails states were determined (Figure 4A). Next, to further investigate auditory versus vestibular specific HC- and SC-differentiation, we calculated cell type specific enrichment scores that were projected onto the *in vitro* data at single cell resolution (Figures 4B,C). Briefly, DEGs for *in vivo* differentiated auditory and vestibular HCs, as well as SCs were calculated using previously published micro array data from postnatal day 1 (P1) mice (Supplemental Figures S2A-D).^45^ Overall, 661 auditory versus vestibular HC-specific DEGs and 846 auditory versus vestibular SC-specific DEGs were determined. Gene set enrichment analysis for the HCLCs and SCLCs was performed using the list of auditory and vestibular HC-specific DEGs and Single-Cell Signature Explorer.^58^ For visualization, auditory and vestibular HC enrichment scores were projected onto the UMAPs (Figures 4B,C). Differential vestibular HC scores were calculated for state S6 (Figure 4B), while auditory HC scores were differentially enriched in state S8 (Figure 4C). At the individual gene level, this finding was supported by *Cib3* being exclusively detected in vestibular like HCLCs, while *Cib2* was found in auditory HCLCs as well as in young vestibular HCLCs.^59^

**Figure 4.**
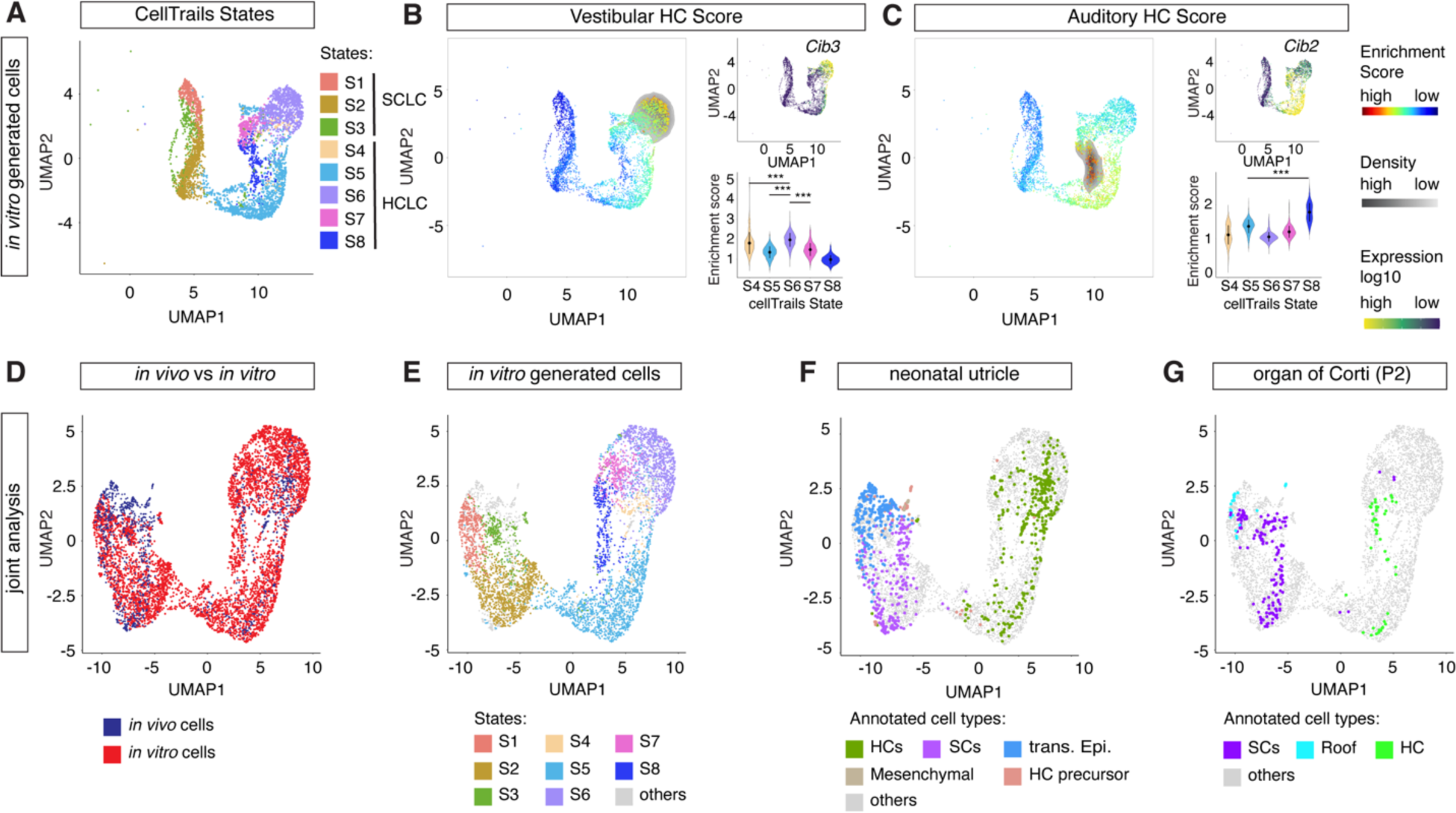
Vestibular and auditory enrichment score analysis. (A-C) Otic epithelial cells were re-clustered using CellTrails to achieve a finer cluster resolution. (A) In total, 3 SCLC clusters and 5 HCLC clusters were identified. (B) Vestibular HC enrichment scores projected onto all otic epithelial cells (left panel). UMAP of all otic epithelial cells with *Cib3* mRNA expression projected (top right panel). Violin plots of vestibular HC enrichment scores for the five HCLC clusters (bottom right panel). CellTrails state ID is color coded. The violin plots show significantly higher vestibular HC enrichment scores in state S6 compared to the remaining states. ***P < 0.001 (Wilcoxon rank sum test). (C) Auditory HC enrichment score analysis with similar representation as in (B). UMAP of all otic epithelial cells with *Cib2* mRNA expression projected (top right panel). The violin plots show significantly higher auditory HC enrichment scores in state S8 compared to the remaining states. ***P < 0.001 (Wilcoxon rank sum test) (bottom right panel). (D-G) Joint analysis of *in vitro* generated HCLCs and SCLCs with primary cells isolated from neonatal mouse utricle and P2 organ of Corti cell types. (D) Joint UMAP projection of all cells with *in vitro* (red) and *in vivo* (blue) origin color coded. (E) UMAP of the joint projection with CellTrails states color coded as outlined in (A) and both *in vivo* data sets color coded in gray. (F) UMAP of the joint projection with neonatal utricle cells annotated as previously published.^7^ Remaining cells color coded in gray. (G) UMAP of the joint projection with P2 organ of Corti cells annotated as previously published.^8^ Remaining cells color coded in gray.

The enrichment analysis did resolve differential scores for vestibular SCs in states S1 and S2 compared to S3, while auditory scores were not significantly different in state S3 (Supplemental Figures S2E,F).

### Joint analysis of *in vitro* generated HCLCs and *in vivo* derived HCs

Bulk based micro array data limit the cell type specific resolution of the enrichment score analysis. To resolve organ specific cell types, we jointly analyzed the transcriptomes of the *in vitro* generated HCLCs and SCLCs together with previously published and annotated scRNA-seq data from neonatal (P2, P4, and P6) utricle^7^ and P2 organ of Corti (Figures 4D-G).^8^ After merging the datasets using Seurat v3^26^ 4,610 *in vitro* derived cells were visualized together with 861 neonatal utricle and 225 P2 organ of Corti cells. First, we assessed the distribution of *in vivo* differentiated cell types in the shared projection (Figure 4D). Cells developed *in vivo* were intermingled with the in HCLC and SCLC clusters and no separation between *in vitro* and *in vivo* developed cells was apparent. As joint analysis was dominated by *in vitro* generated HCLCs and SCLCs, the overall topology of the UMAP projection was preserved compared to projecting *in vitro* derived cells alone (Figure 5E). The mutually exclusive distribution of *in vivo* derived utricle and organ of Corti HCs (Figures 4F,G) confirmed the asymmetric distribution of micro-array based organ specific enrichment scores (Figures 4B,C). Together, co-analysis of our organoid derived cells with previously published data of *in vivo* developed HCs and SCs indicate differentiation of cochlear and vestibular like cell types *in vitro*.

**Figure 5.**
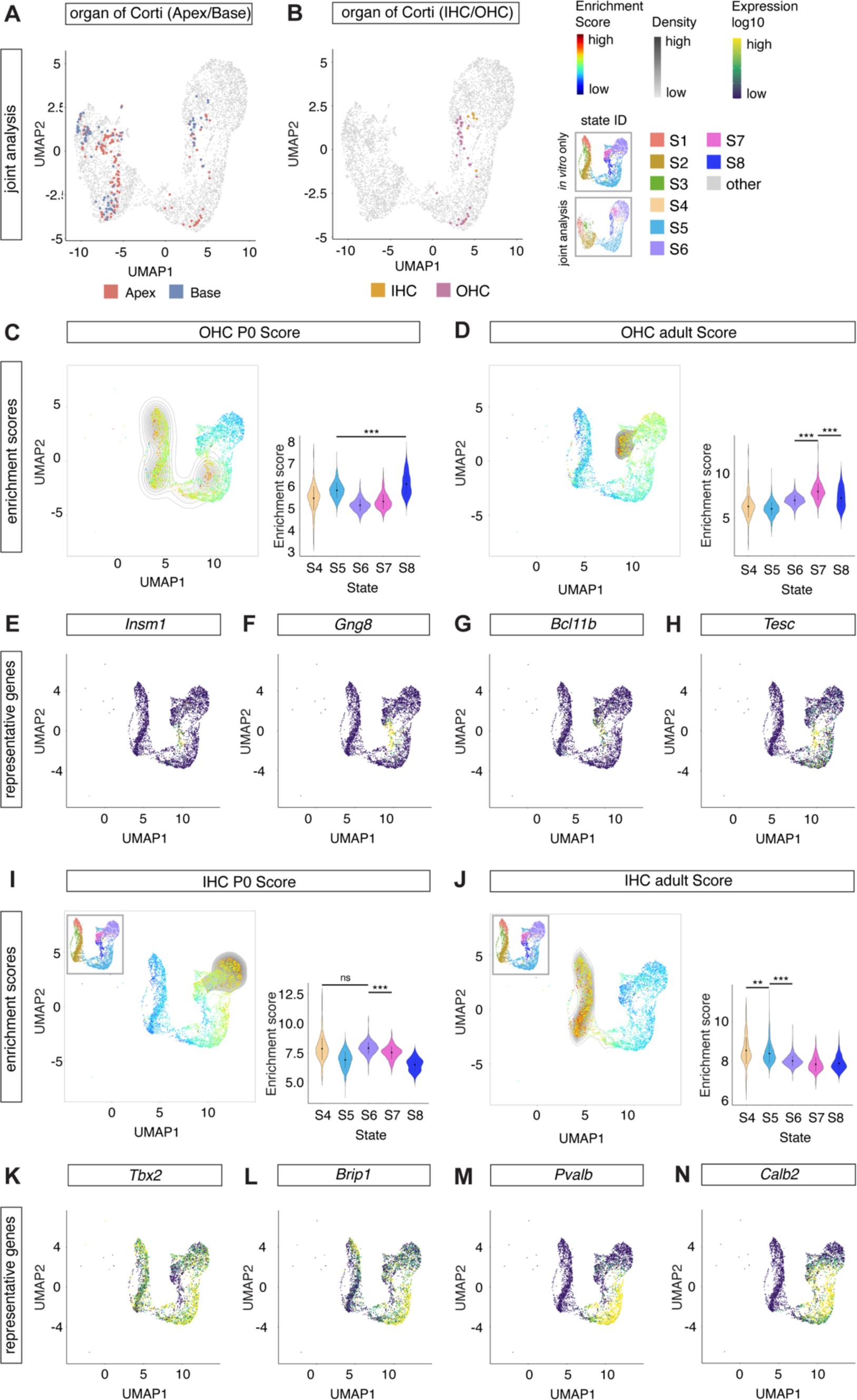
Postnatal and adult IHC and OHC enrichment score analysis. (A,B) Joint analysis of *in vitro* generated HCLCs and SCLCs with primary cells isolated from neonatal mouse utricle and P2 organ of Corti cell types. (A) Joint UMAP projection of all cells with apical (red) and basal organ of Corti cells (blue) color coded. Remaining cells color coded in gray. (B) Joint UMAP projection of all cells with IHCs (gold) and OHCs (purple) color coded. Remaining cells color coded in gray. (C-N) IHC vs OHC enrichment score analysis. (C) P0 OHC enrichment scores projected onto all otic epithelial cells (left panel). Violin plots of P0 OHC enrichment scores for the five HCLC clusters (right panel). CellTrails state ID is color coded. The violin plots show significantly higher P0 OHC enrichment scores in state S8 compared to the remaining states. ***P < 0.001 (Wilcoxon rank sum test). (D) Adult OHC enrichment score analysis with similar representation as in (C). The violin plots show significantly higher adult OHC enrichment scores in state S7 compared to the remaining states. ***P < 0.001 (Wilcoxon rank sum test). (E-H) UMAP of all otic epithelial cells with OHC markers *Insm1* (E), *Gng8* (F), *Bcl11b* (G), *and Tesc* (H) mRNA expression projected. (I,J) P0 (I) and adult (J) IHC enrichment score analysis with similar representation as in (C,D). The violin plots show significantly higher P0 IHC enrichment scores in states S4 and S6 compared to the remaining states. ***P < 0.001 (Wilcoxon rank sum test) (I, right panel). In comparison, no statistically significant enrichment was fount for the adult IHC scores in the HCLC compartment (J, right panel). (K-N) UMAP of all otic epithelial cells with IHC markers *Tbx2* (K), *Brip1* (L), *Pvalb* (M), *and Calb2* (N) mRNA expression projected.

### Differentiation of organ of Corti like HCLCs *in vitro*

To further analyze organ specific characteristics, we revisited the joint alignment analysis of *in vitro* and *in vivo* generated inner ear cell types (Figure 5A). Differences along UMAP2 likely correspond to differences in developmental maturation stages as visualized by RNA velocity (Figure 3A). Therefore, we plotted the anatomical origin, apex versus base, for the organ of Corti derived HCs (Figure 5A). At P2, basal cochlear HCs are more mature compared to their apical counterparts. Hence, more mature HCs isolated from the base clustered with state S8 and immature HCs isolated from the apex localized to the immature *in vitro* derived cells as represented by state S5.

While development and maturation proceed in gradients extending from the base towards the apex of the cochlea, morphogens such as SHH, BMP7, and RA^60–62^ pattern regional identities of cochlear floor cells depending on their relative position along the longitudinal (tonotopic) axis of the cochlea.^63^ During embryonic development, regional identity is mirrored in tonotopic gene expression pattern of genes such as *Fst*, *Hmga2*, and *A2m.*^62, 64, 65^ However, during organoid development, none of the candidate genes was detected in HCLCs nor in SCLCs (Supplemental Figure S3A). Together, these findings suggest that no regional identity is mediated by the self-guiding protocol.

To further analyze organ specific characteristics, we revisited the joint alignment analysis of *in vitro* and *in vivo* generated inner ear cell types and projected organ of Corti inner HC (IHC) and outer HC (OHC) identities (Figure 5B). Both, P2 IHCs and OHCs were projected intermingled and spread along UMAP2. Next, enrichment scores for perinatal and adult IHCs and OHCs were calculated using bulk RNA-seq data (Figures 5C-D, I-J).^66^ Perinatal OHC scores were enriched in *in vitro* generated cluster S8, while adult OHC scores were differentially higher in the more mature HCLCs of cluster S7 (Figures 5C,D). This finding was supported by OHC specific genes such as *Insm1*, *Gng8*, *Bcl11b*, and *Tesc*^66, 67^ being differentially expressed in cluster S8 (Figures 5E-H). In comparison, perinatal IHCs scores were enriched in states S4 and S6, while adult IHC scores were not enriched in the HCLCs (Figures 5E,F). Interestingly, both states S4 and S6 were also characterized by differential vestibular enrichment scores (Figure 4B), a finding that is likely due to a high number of shared genes between vestibular and IHCs such as *Tbx2*^67, 68^, *Brip1*^66, 69^, *Pvalb*^70, 71^, and *Calb2*^5, 72^ (Figures 5K-N and Supplemental Figures S2A-D). In summary, the results presented above indicate that a small proportion of the HCLCs followed an OHC trajectory with maturation beyond the perinatal stage as indicated by the adult OHC enrichment scores. Maturation of young IHC like HCLCs either does not occur or is obscured by vestibular like differentiation. A progression towards more mature IHC like stages comparable to the OHC like HCLCs was not observed.

### Differentiation of vestibular like HCLCs *in vitro*

Our initial analysis and previous reports^18, 73–75^ indicate that the majority of *in vitro* generated HCLCs differentiate into vestibular phenotypes by default. To elucidate the phenotypic variation of the vestibular like HCLCs, we calculated enrichment scores for perinatal extrastriolar Type II, extrastriolar Type I, and striolar Type I HCs based on previously published data ^7^ using Single-Cell Signature Explorer^58^ (Figures 6A-C). Cells of the state S5 were characterized by differential Type II extrastriolar HC scores and differential expression of the Type II marker gene *Anxa4*^5^ was detected. Extrastriolar Type I HC scores were enriched in state S6 and the Type I HC marker *Spp1*^5^ was differentially expressed in state S6 as well. Finally, state S4 exhibited differential striolar Type I HC scores although at lower levels. Similarly, *Ocm*, a marker for striolar HCs,^6^ was differentially expressed in state S4.

**Figure 6.**
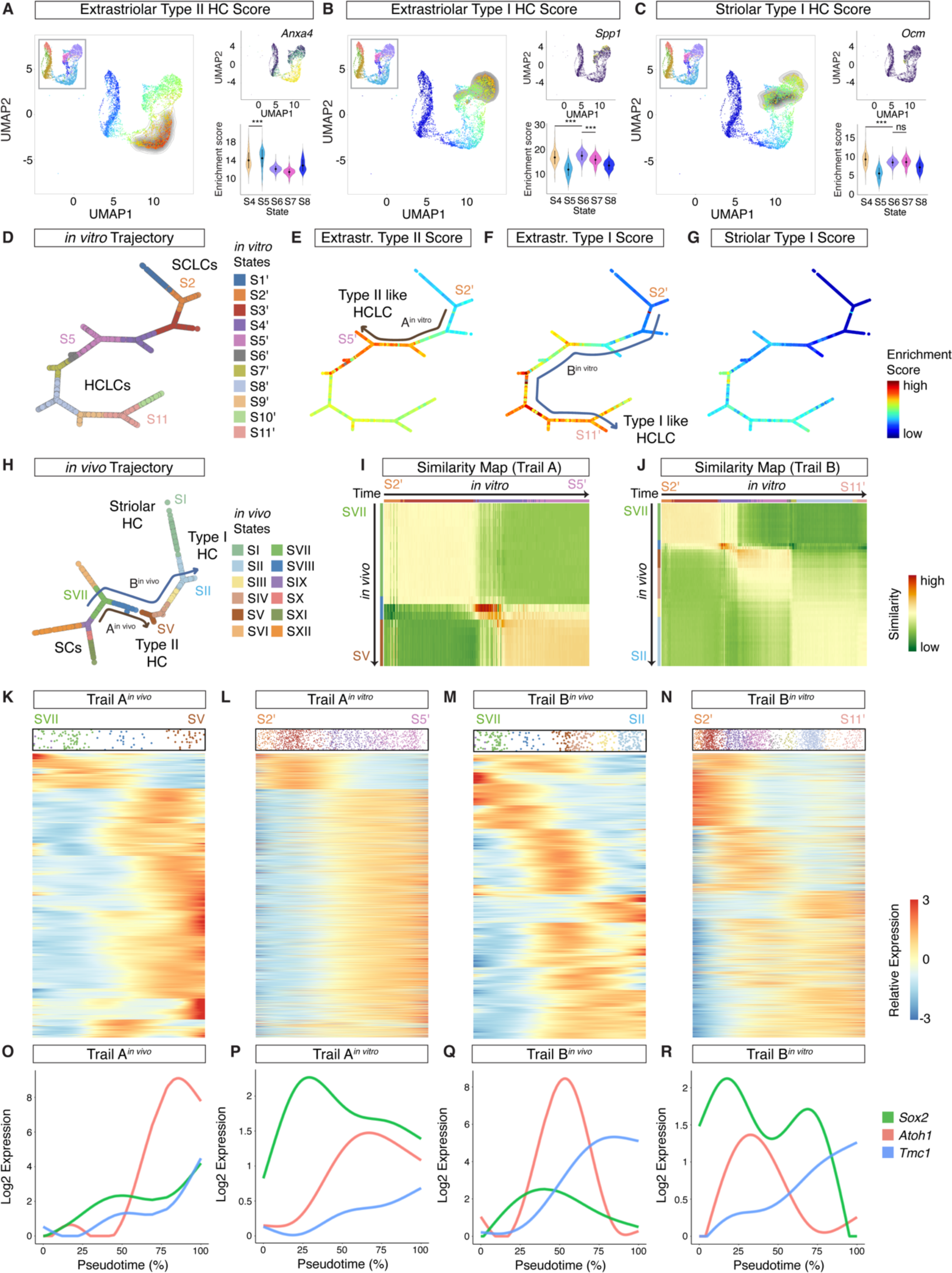
Comparative trajectory analysis of vestibular HCs and HCLCs. (A) Extrastriolar Type II HC enrichment scores projected onto all otic epithelial cells (left panel). The inset shows similar UMAP with CellTrails states color coded for orientation. UMAP of all otic epithelial cells with *Anxa4* mRNA expression projected (top right panel). Violin plots of extrastriolar Type II HC enrichment scores for the five HCLC clusters (bottom right panel). The violin plots show significant enrichment of extrastriolar Type II HC enrichment scores in state S5 compared to the remaining states. ***P < 0.001 (Wilcoxon rank sum test). (B) Extrastriolar Type I HC enrichment score analysis with similar representation as in (A). UMAP of all otic epithelial cells with *Spp1* mRNA expression projected (top right panel). The violin plots show significantly higher extrastriolar Type I HC enrichment scores in state S6 compared to the remaining states. ***P < 0.001 (Wilcoxon rank sum test) (bottom right panel). (C) Striolar Type I HC enrichment score analysis with similar representation as in (A). UMAP of all otic epithelial cells with *Ocm* mRNA expression projected (top right panel). The violin plots show significant enrichment of the striolar Type I HC scores in state S4 compared to the remaining states. ***P < 0.001 (Wilcoxon rank sum test) (bottom right panel). (D) Inferred branching trajectory graph for *in vitro* generated SCLCs and HCLCs with 11 states (S’). Organ of Corti like HCLCs were excluded. SCLC and HCLC identities are indicated. (E-G) Extrastriolar Type II (E), extrastriolar Type I (F), and striolar Type I (G) enrichment scores projected onto the *in vitro* trajectory. Enrichment scores are color coded for high (red) and low (blue). Inferred extrastriolar Type II (trail A*^in vitro^*) and extrastrolar Type I (trail B*^in vitro^*) are indicated. (H) Inferred branching trajectory graph for *in vivo* developed SCs and HCs with 12 states as previously published.^7^ CellTrails states are color coded. Cell type annotations and inferred extrastriolar Type II (trail A*^in vivo^*) and extrastrolar Type I (trail B*^in vivo^*) trajectories are indicated. (I) Cosine similarity matrix comparing trail A*^in vivo^* (y-axis) with trail A*^in vitro^* (x-axis). Cells are ordered along trail A and aggregated into metacells. Similarity is color coded from high (red) to low (green). (J) Cosine similarity matrix comparing trail B*^in vivo^* (y-axis) with trail B*^in vitro^* (x-axis). Similar data representation as in (I). (K) Dynamic gene expression for 3730 genes along trail A*^in vivo^*. The cell density of the trail is indicated at the top of the heatmap. The order of the genes along the y-axis was determined by the similarity of their expression patterns. (L) Dynamic gene expression along trail A*^in vitro^*. Order of the genes and data representation are similar to (K). (M) Dynamic gene expression for 6890 genes along trail B*^in vivo^*. The order of the genes along the y-axis was determined by the similarity of their expression patterns. (N) Dynamic gene expression along trail B*^in vitro^*. Order of the genes and data representation are similar to (M). (O-R) Expression dynamics of *Sox2* (green), *Atoh1* (red), and *Tmc1* (blue) along trail A*^in vivo^* (O), trail A*^in vitro^* (P), trail B*^in vivo^* (Q), and trail B*^in vitro^* (R).

RNA velocity as well as the gene expression pattern of individual genes such as *Atoh1* and *Tmc1* suggest that the cells collected represent different stages of HC maturation. To further investigate the gene expression dynamics of vestibular HCLC development, single cell trajectory analysis was performed using CellTrails.^57^ After excluding state S8 organ of Corti like HCs, eleven states (S’) were identified (Figure 6D). Projecting the cell trails states onto the trajectory, SCLCs (S1’-S3’) and HCLCs (S4’-S11’) were located at opposite ends of the trajectory. Next, we projected Type I and II HC specific enrichment scores onto the reconstructed trajectory to identify cell type specific trajectories. The extrastriolar Type II HC trajectory (trail A*^in vitro^*) was defined as a sub-trail starting from the S2’ branching point between sensory SCLCs and NSECs towards state S5’ HCLCs (S2’ → S3’ → S4’→ S5’) (Figure 6E). Similarly, the extrastriolar Type I trajectory (trail B*^in vitro^*) started at the S2’ branching point but extended further to an end point constituted by state S11’ HCLCs (S2’ → S3’ → S4’→ S5’→ S6’ → S7’ → S8’ → S9’→ S11’) (Figure 6F). Considering the low scores for the striolar Type I HCLCs, we refrained from defining a designated trajectory (Figure 6G).

*In vivo* versions of the Type I and Type II HC trajectories (trails A*^in vivo^* and B*^in vivo^*) reconstructed from perinatal utricle cells have previously been published (Figure 6H).^7^ To analyze the similarity of the gene expression dynamics during *in vitro* and *in vivo* HC development, we aligned the trajectories in a Cosine similarity matrix and visualized the results using a heatmap (Figures 6I,J). We denoised the data by aggregating the six nearest neighboring cells of the trajectory into metacells. Metacells from the *in vitro* generated SCLCs (S2’) aligned to *in vivo* developed SCs (SVII), Type II like HCLC metacells (S5’) aligned to perinatal Type II HCs (SV) (Figure 6I), and Type I like HCLCs (S11’) aligned to perinatal Type I HC metacells (SII) (Figure 6J). The highest similarity scores were arranged in a diagonal across the heatmaps, indicating that start and end points for the *in vivo* and *in vitro* derived trajectories were aligned properly. Highest similarity scores were characteristic for early HC precursors of states S4’ and SVIII.

To compare gene expression dynamics along the individual trajectories at whole transcriptome level, we fitted generalized additive models of gene expression as a function of pseudotime using CellTrails as outlined by Taha et al. (Figures 6K-N). This allowed us to systematically identify genes that are changing along a specific trail to determine similarities and differences between *in vivo* and *in vitro* development. In case of the Type II trajectory (trail A), 3,730 genes were dynamically changing along the trajectory as previously published (Figure 6K). We used the order of genes in the heatmap as a template to analyze temporal gene expression along the Type II *in vitro* trajectory (Figure 6L). Overall, 3,491 genes (93.6%) were found with similar expression dynamics compared to the *in vivo* trail. The remaining genes were not detected in the *in vitro* dataset. Similarly, 6890 genes in the Type I trajectory (trail B) followed similar dynamics in expression, while 641 of the genes found with differential expression along the *in vivo* trail were not detected *in vitro* (Figures 6M,N). Aiming to elucidate differences between *in vitro* and *in vivo* HC differentiation, genes that were absent in the *in vitro* trajectory were extracted and absolute expression values were plotted along the *in vivo* trajectory (Supplemental Figure S4A). Based on this analysis, most of the candidate genes were expressed close to the limits of detection in the smart-seq based utricle data set. A GO-term analysis of the same genes revealed significant enrichment for 5 GO terms, potentially indicating deficits in synapse formation for the *in vitro* generated HCs (Supplemental Figure S4B).

To visualize differentiation of HCLCs into Type I and Type II like fates, we focused on the expression of three candidate genes (Figures 6 O-R). Sox2 is a transcription factor that is expressed in sensory SCs and Type II HCs. Confirming the expression of the Type II HC specific marker, continuous *Sox2* expression was found for the *in vivo* and *in vitro* versions of trail A (Figures 6O,P). *Atoh1*, the master regulator of HC fate,^29^ is induced after Sox2 upregulation in trail A for both *in vivo* and *in vitro* generated HCs. Similarly, *Tmc1*, associated with the function of HCs,^56^ exhibits a steady increase in expression during the time course reconstructed by trail 1 for both paradigms. Type I HCs in comparison do not express significant levels of *Sox2*. However, a transient expression of the transcription factors was detected for both *in vivo* and *in vitro* differentiated Type I HCs (Figures 6Q – R). Similarly, the *Atoh1* expression pattern is of transient nature and *Tmc1* shows a steady increase in expression comparable to what has been observed in Type II HCs *in vivo* and *in vitro*.

Together, these findings demonstrate that *in vitro*-based HCLC differentiation is not restricted to a small number of genes such as *Atoh1* and *Sox2*, but closely resembles gene expression pattern and dynamics as seen *in vivo* at whole transcriptome level.

## Discussion

Despite the enormous potential that has been attributed to inner ear organoids in regenerative medicine and basic science research, our understanding of how HCLCs develop *in vitro* and the knowledge about their phenotypic diversity remains limited. In this study we used scRNA-seq to transcriptionally profile *in vitro* generated HCLCs and SCLCs and compare the data to different auditory and vestibular cell types. We studied so called late-stage organoids to elucidate the differentiation potential and we aimed to determine the level of maturation present in *in vitro* differentiated cell types.

### Organ Specification

Early stages of organoid differentiation—from the pluripotent stem cell to the otocyst stage— are highly controlled with timed chemical cues that recapitulate major signals *in vivo*. In contrast, in most current organoid culture protocols, later stages of organoid differentiation—from the otocyst stage to the functional HCLC—are self-guided with little outside influence. Following the Koehler^19^ protocol, we cultured organoids between 16 and 21 DIV to generate SCLCs and HCLCs. Based on a combination of trajectory-based analysis using RNA velocity, auditory and vestibular enrichment score analysis, and joint projection with *in vivo* developed HCs, a separation into auditory and vestibular HCLCs was apparent. In comparison, even from the earliest development of *in vitro* protocols, inner ear organoid HCs appear to express morphological and biochemical characteristics like those in vestibular organs.^18, 73^ For example, in a large-scale electrophysiological screen of 153 mouse organoid derived HCs, the majority exhibited mechanotransduction currents and basolateral conductance most similar to Type II utricular HCs.^76^ More recently, single-cell transcriptomic evaluations of organoids from human pluripotent stem cells support the general notion that organoid HCLCs default to a vestibular like fate.^74, 75^ Even so, at least one report has shown evidence of both vestibular and cochlear HC fates with only slight modifications to the original Koehler protocol^77^ and modulations of HH and WNT signaling were deemed sufficient to tip the balance towards cochlear cell fates.^24^ Using scRNA-seq, we profiled a total of 3,029 HCLCs and in an unbiased clustering approach we determined that a significant number of HCLCs acquired a transcriptional signature reminiscent of auditory HCs. It remains to be determined if those cells develop in designated cysts or organoids that exclusively give rise to auditory HCLCs or if they develop intermingled with their vestibular counter parts. Interestingly, virtually all the cells exhibiting statistically significant organ of Corti enrichment scores, namely the cells in CellTrails state S8, developed into OHC like HCs. Also, a progression from postnatal to adult OHC gene expression pattern was evident based on gene set enrichment scores. However, cardinal genes associated with functionally mature OHCs, such as *Slc26a5* encoding for PRESTIN and *Kcnq4*^78, 79^ were notably absent in the organ of Corti like HCLC population raising the question if prolonged culture duration would be sufficient to allow further maturation. In comparison, highest perinatal IHC enrichment scores were found in cells that also share high similarity with extrastriolar Type I like HCs, and there was no evidence for a progression towards maturation of adult like IHC like cells. This finding lends further support to the hypothesis that the OHC fate represents a default state in auditory HC development as it has been postulated based on the role of the transcription factor TBX2 in IHC development.^67^ TBX2 was identified as a master regulator for the induction of IHCs that would otherwise develop into OHCs in absence of the transcription factor. Hence, absence of *Tbx2* in HCLCs developing into OHC like cells was confirmed; however, cells of the vestibular trajectory were found to express *Tbx2*. This finding suggests, although *Tbx2* was identified as master regulator of IHC fate, expression of *Tbx2* in organoid derived HCLCs is not sufficient to further drive IHC maturation.

The majority of HCLCs develop into vestibular like phenotypes. Based on previous reports, this finding was expected; however, the accuracy of how HCLCs recapitulate *in vivo* HC development was intriguing. While *in vivo* differentiation from HC precursors into Type II vestibular HCs appears like a binary switch in gene expression pattern, the differentiation into Type I vestibular HCs requires a more intricate temporal coordination of transient gene expression.^7^ For both organoid trajectories of differentiation into Type I and Type II HCLCs, a resemblance of the gene expression dynamics at whole transcriptome level was apparent. Moreover, the similarity map comparing *in vivo* and *in vitro* differentiation highlights two putative bottlenecks. First, the transition from SC to HC precursor is characterized by highest similarity between *in vivo* and organoid based gene expression. This finding could indicate that a defined set of genes is necessary to mediate the progression from SC like progenitor into HC precursor, while other sections of the trajectory tolerate higher variability. A second bottleneck was related to state S6’, an intermediate state of the Type I specific trajectory (Trail B). The small number of cells contributing to state S6 compared to the other states could indicate that this state is not stable and transition proceeds fast. However, further characterization and biological relevance of state S5’ remains to be determined.

With respect to SCLCs, a segregation into auditory or vestibular like cell types was less apparent. Overall, SCLCs resemble vestibular gene expression signatures based on enrichment scores; however, differentiation into a more generic supporting cell type expressing less organ specific gene sets could serve as alternate explanation.

Even though, our study focused on so called late-stage organoids collected between 16 and 21 DIV, the UMAP representation visualized a continuum between SCLCs and HCLCs that allowed for the reconstruction of a developmental trajectory similar to what has been described for *in vivo* utricle development.^7^ This finding suggests protracted development of vestibular HCLCs from the SCLC pool; although, a designated population of stem cell like cells replenishing the SCLCs was not identified. Analysis of a more expanded culture range, both younger and older, may be required to identify common progenitors as well as the extent of maturation in this culture system.

Differences in *in vivo* and *in vitro* differentiated HCs were attributed to biological and technological reasons. 1) We found differential GO terms indicating biological deficits in synapse formation for the HCLCs. While *in vitro* synaptogenesis for inner ear organoids was reported previously,^20, 80^ the synaptic structures appear incomplete and atypical for neonatal vestibular HCs. Synapse formation, however, regulates hair cell excitability and maturation, and aberrant synapse development leads to the maintenance of immature electrical features in vestibular HCs.^81^ 2) Further, the different sequencing platforms applied (10x Genomics vs smart seq2) are associated with different limits of detection, which in turn might explain why a number of low abundance genes was not detected in the *in vitro* data set.

### Regional Specification

HCs not only differentiate into organ specific subtypes, but also regional specification shapes the function of the cochlear and vestibular HCs. For example, molecular and physiological differences along the tonotopic axis shape the function of auditory HCs and are critical in frequency specific hearing.^10^ Morphogens such as SHH, RA, and Bmp7 have been identified to pattern the apex-to-base axis in the mouse and chicken cochlea.^60–62^ Naturally, regional specification occurs after organ specification; hence, current versions of the organoid protocol do not provide specific guidance in this respect. Our integrated analysis of *in vivo* developed HCs with the HCLCs indicates that the cochlear trajectory, mainly represented by OHC like HCLCs, resembles an apex-to-base order. However, absence of tonotopic marker genes such as *Fst*^65^, suggests that the OHC like trajectory reflects developmental differences rather than spatial identity. Moreover, absence of tonotopic marker genes suggests that HCLCs do not acquire apical nor basal phenotypes by default.

Vestibular HCs in comparison, exhibit distinctly different neural activity between striolar and extrastriolar regions.^6, 82^ Like in patterning of the tonotopic axis of the cochlea, RA has been identified to play a role in patterning the striolar-extrastriolar axis of the mouse vestibular organs with extrastriolar identity being associated with higher RA levels compared to the striolar identity. Interestingly, the transcriptional profiles of the vestibular like HCLCs indicate differentiation of extrastriolar Type I and II like HCs, suggesting varied RA signaling between aggregates or within individual organoids. Neither RA itself, nor its precursor retinol (vitamin A) are supplemented with the cell culture media during generation of the organoids. Although the chemical composition of media components throughout the culture protocol are highly defined, the addition of animal-derived extracellular matrix Matrigel with notorious lot-to-lot variability raises the possibility of biologically relevant retinoids within the culture paradigm. Given the largely extrastriolar identity of our organoid HCLCs, we conclude that either undefined media components provide exogenous retinoids and induce regional specification or additional signaling pathways are involved in striolar-extrastriolar patterning.

In summary, the results of this study indicate that without further guidance HCLCs differentiate into vestibular and auditory phenotypes randomly, although the extrastriolar vestibular trajectory appears to be preferred. Future efforts in modulating different signaling pathways following the otocyst stage *in vitro* hold the promise to control organ as well as region specific differentiation using the organoid model.

---

**Supplemental Figure S1.**
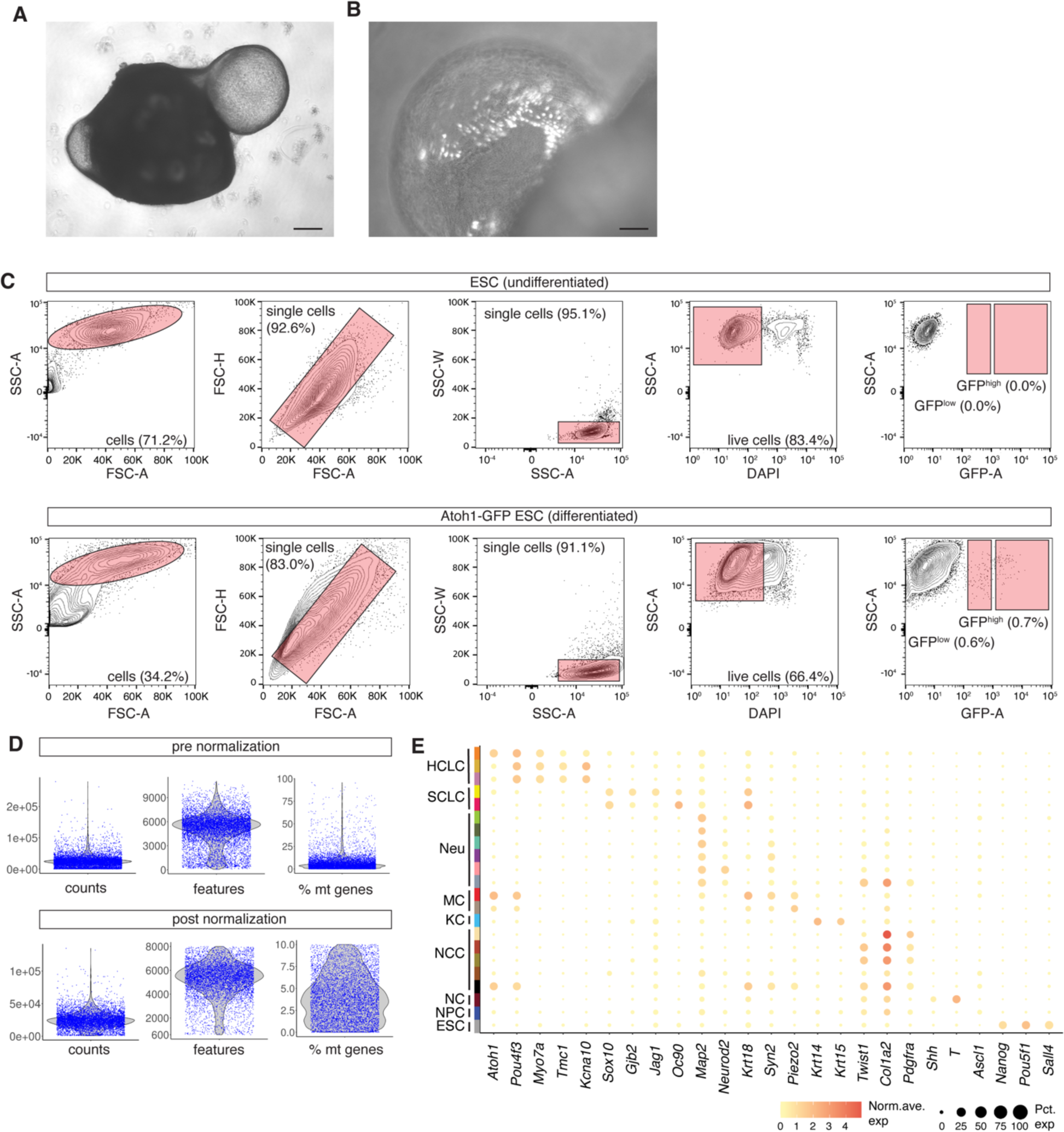
Sample preparation and transcriptional profiling. (A) Phase-contrast image of 20 DIV aggregate with protruding cystic organoid. (B) High-magnification phase-contrast image of cystic organoid overlayed with GFP fluorescence. (C) FACS plot and gating strategy to isolate cells expressing GFP at high and low levels. 1^st^ row: undifferentiated ESCs not expressing GFP as a negative control. 2^nd^ row: experimental *Atoh1*-GFP ESCs after going through the differentiation protocol. (D) Raw data before and after quality control. (E) Dot plot of candidate gene expression levels for each scRNA-seq cluster. The dot size represents the percentage of cells expressing a given transcript for the clusters. Gene expression levels are color coded from yellow (low) to red (high). HCLC – HC like cell, SCLC – SC like cell, Neu – neuronal like cell, MC – Merkel like cell, KC – keratinocyte like cell, NCC – neural crest like cell, NC – notochord like cells, NPC – neuronal precursor like cell, ESC – embryonic stem cell. Scale bars: 100 µm in A and 25 µm in B.

**Supplemental Figure S2.**
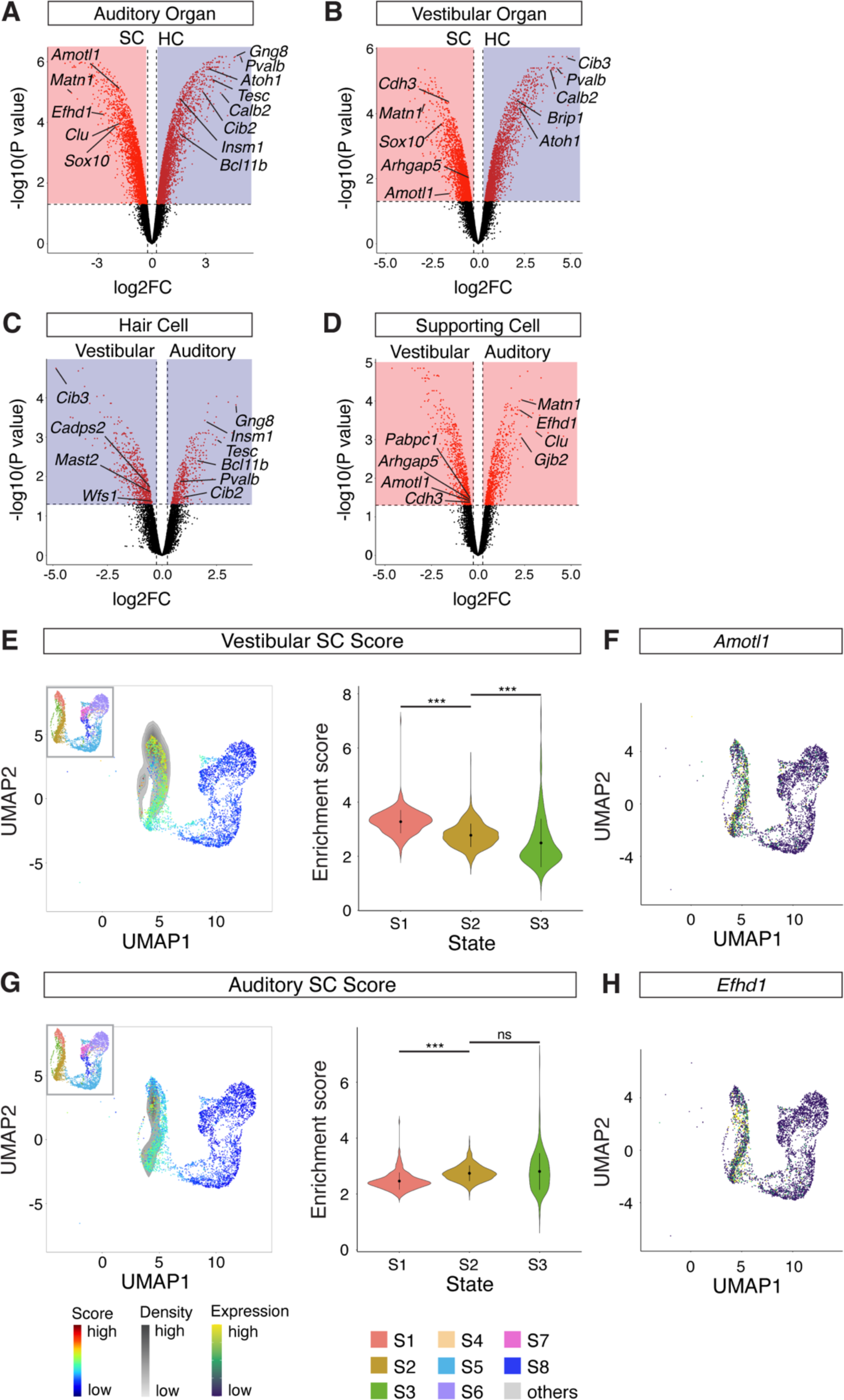
Auditory and vestibular enrichment score analysis. (A) Volcano plots of DEGs comparing P2 organ of Corti SCs (red) and HCs (blue). Cutoff: adjusted p-value < 0.05 and absolute value of log2FC > 0.25. (B) Volcano plots of DEGs comparing P2 vestibular SCs (red) and HCs (blue). Cutoff: adjusted p-value < 0.05 and absolute value of log2FC > 0.25. (C) Volcano plots of DEGs comparing P2 vestibular HCs (blue) and auditory HCs (blue). Cutoff: adjusted p-value < 0.05 and absolute value of log2FC > 0.25. (D) Volcano plots of DEGs comparing P2 vestibular SCs (blue) and auditory SCs (blue). Cutoff: adjusted p-value < 0.05 and absolute value of log2FC > 0.25. The auditory HC enrichment score was calculated using a gene set that was differentially expressed in auditory HCs compared to vestibular HCs as well as differentially expressed in auditory HCs compared to auditory SCs. Using a similar strategy, we calculated vestibular HC, auditory SC, and vestibular SC enrichment scores. (E) Vestibular SC enrichment scores projected onto all otic epithelial cells (left panel). The inset shows the similar UMAP with CellTrails states color coded for orientation. Violin plots of vestibular SC enrichment scores for the three SCLC clusters (right panel). The violin plots show significant enrichment of vestibular SC enrichment scores in state S1 compared to the remaining states. ***P < 0.001 (Wilcoxon rank sum test). (F) UMAP of all otic epithelial cells with *Amotl1* mRNA expression projected. (G) Auditory SC enrichment scores projected onto all otic epithelial cells (left panel). Violin plots of auditory SC enrichment scores for the three SCLC clusters (right panel). The violin plots show significant enrichment of auditory SC enrichment scores in state S2 compared to the state S1 but not compared to S3. ***P < 0.001 (Wilcoxon rank sum test). (H) UMAP of all otic epithelial cells with *Efhd1* mRNA expression projected.

**Supplemental Figure S3.**
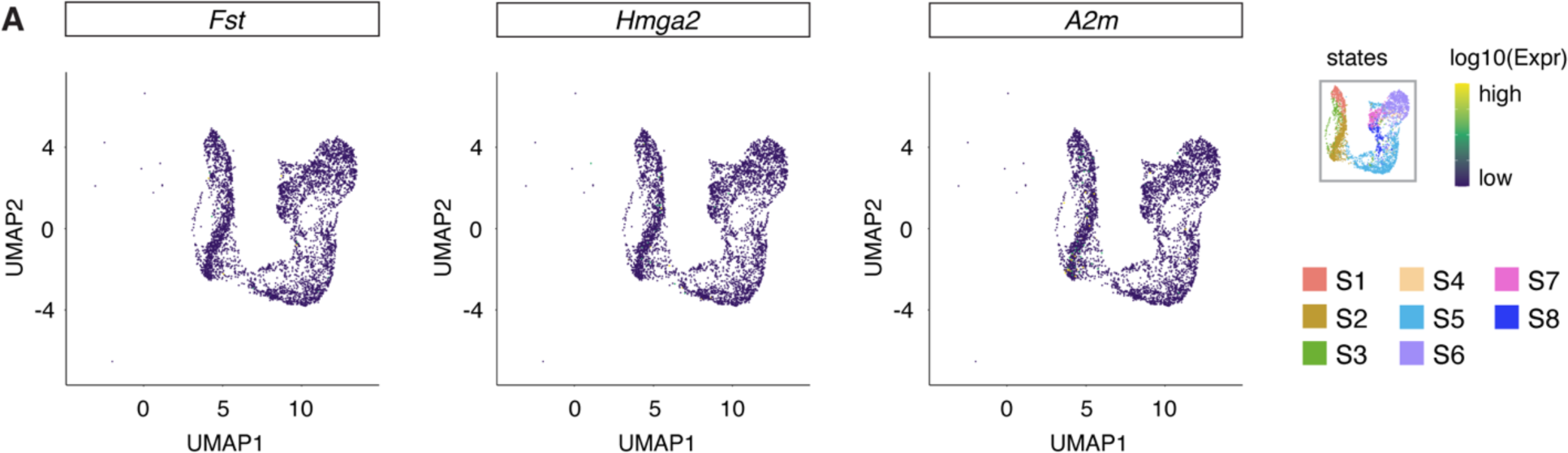
Expression pattern of tonotopic marker genes. (A) UMAP of all otic epithelial cells with tonotopic marker *Fst*, *Hmga2*, *and A2m* mRNA expression projected.

**Supplemental Figure S4.**
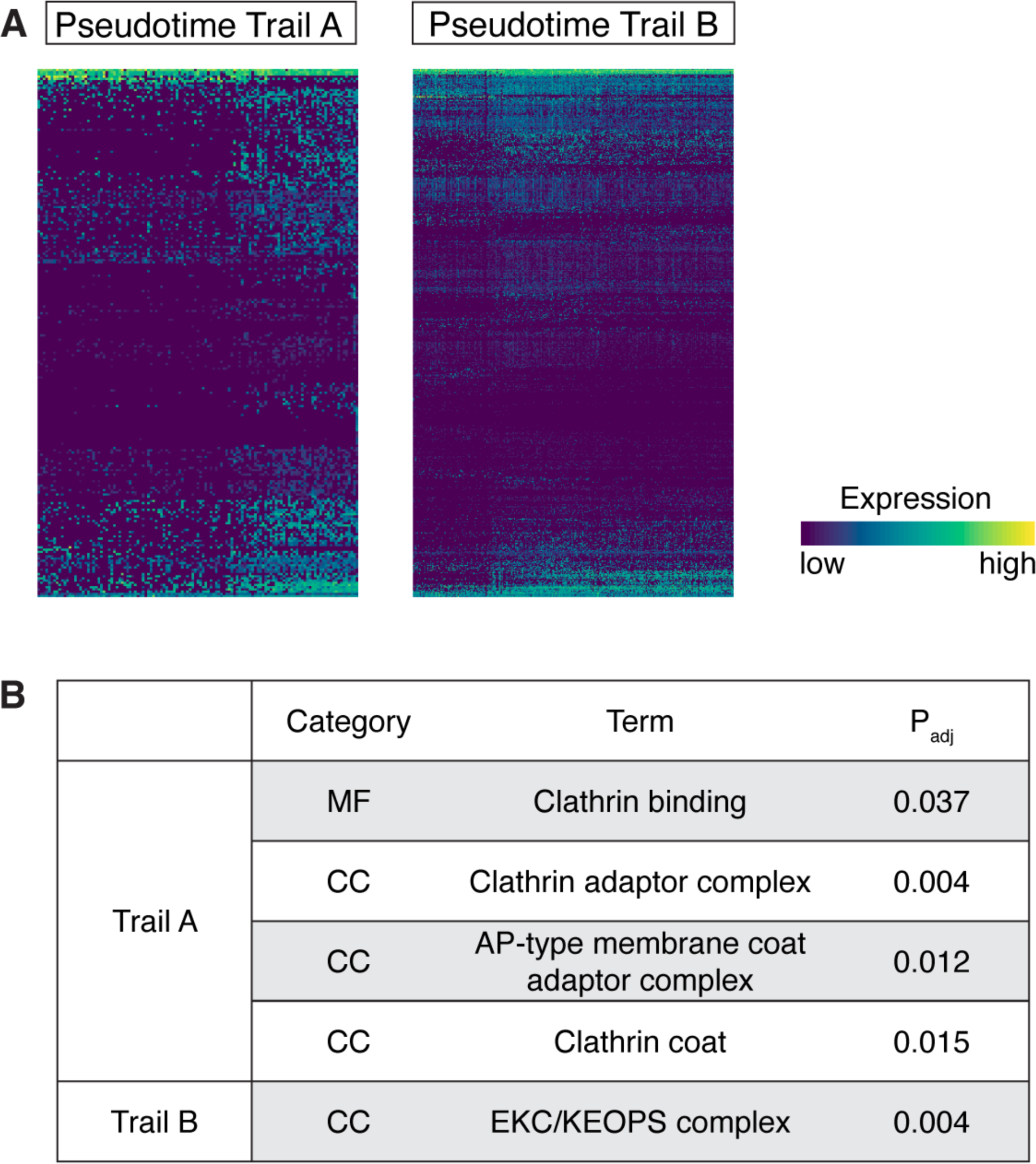
Differences between *in vitro* and *in vivo* differentiation along the extrastriolar Type I and II trajectories. (A) Dynamic gene expression for 239 genes along trail A*^in vivo^*. Gene candidates plotted in the heatmap were detected in the *in vivo* data set but not *in vitro* (left panel). (A) Dynamic gene expression for 641 genes along trail B*^in vivo^* (right panel). All candidate gene were detected *in vivo* but not *in vitro*. (B) GO term analysis for the gene sets that were detected during *in vivo* development only.

## Methods

### Key resources table

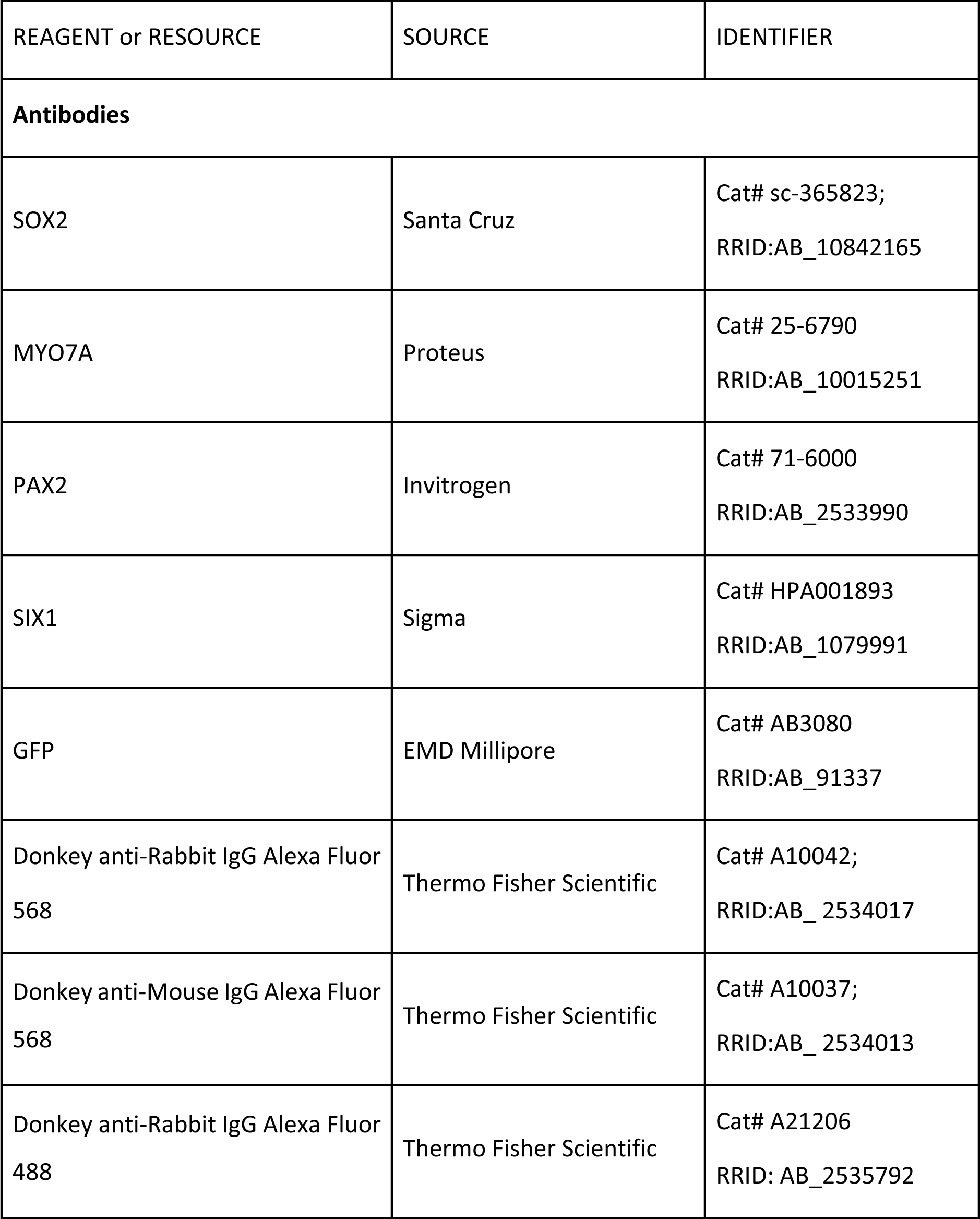
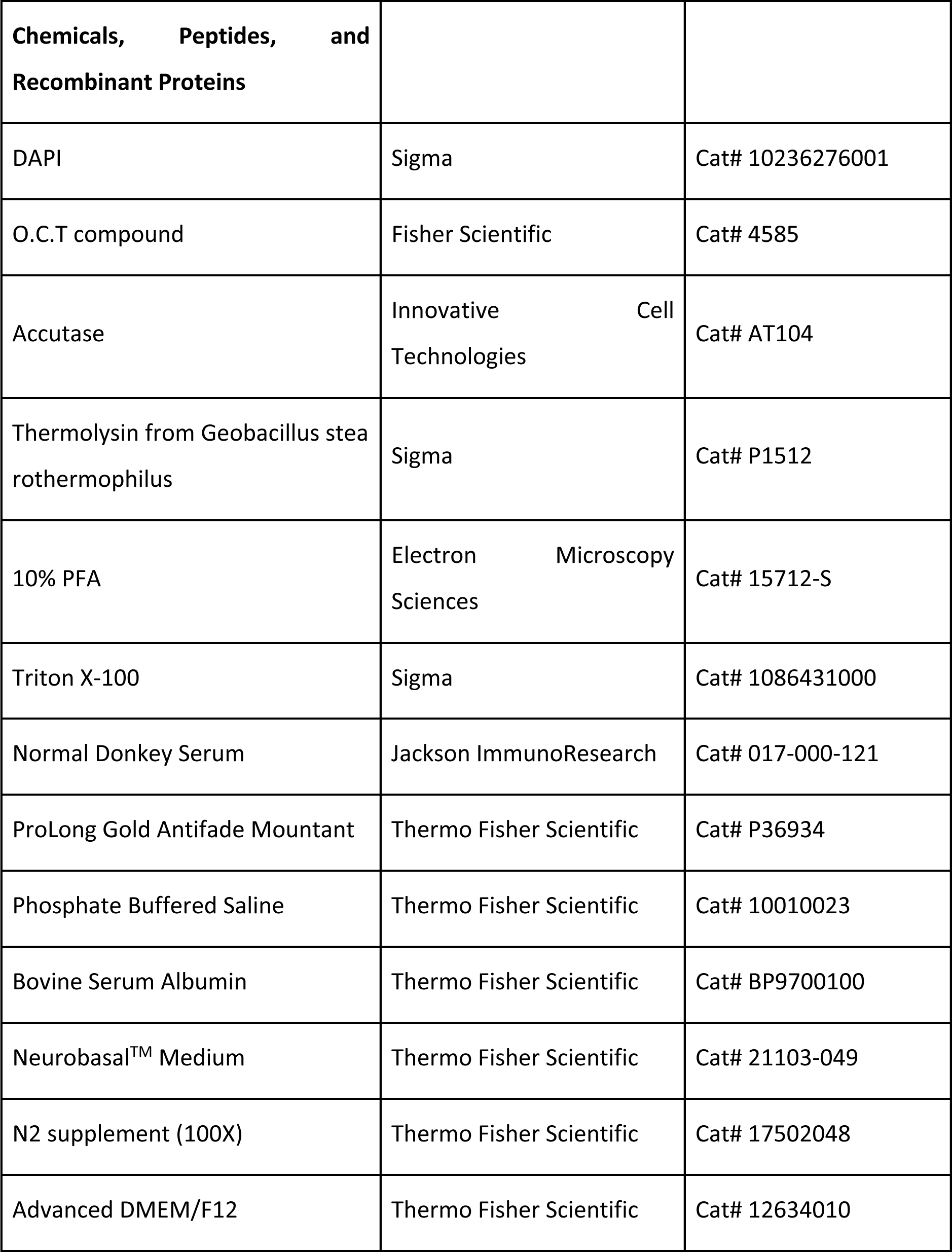
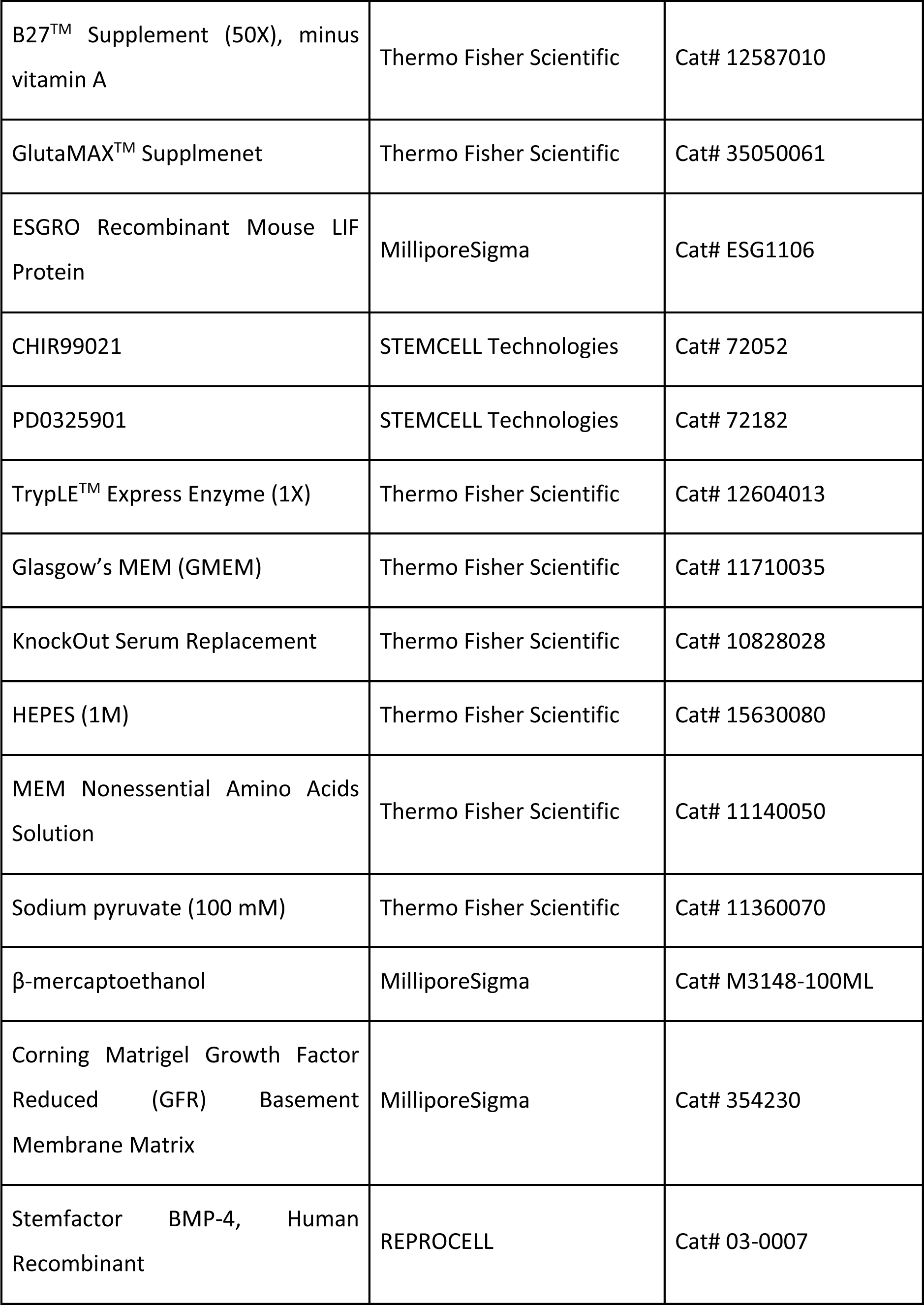
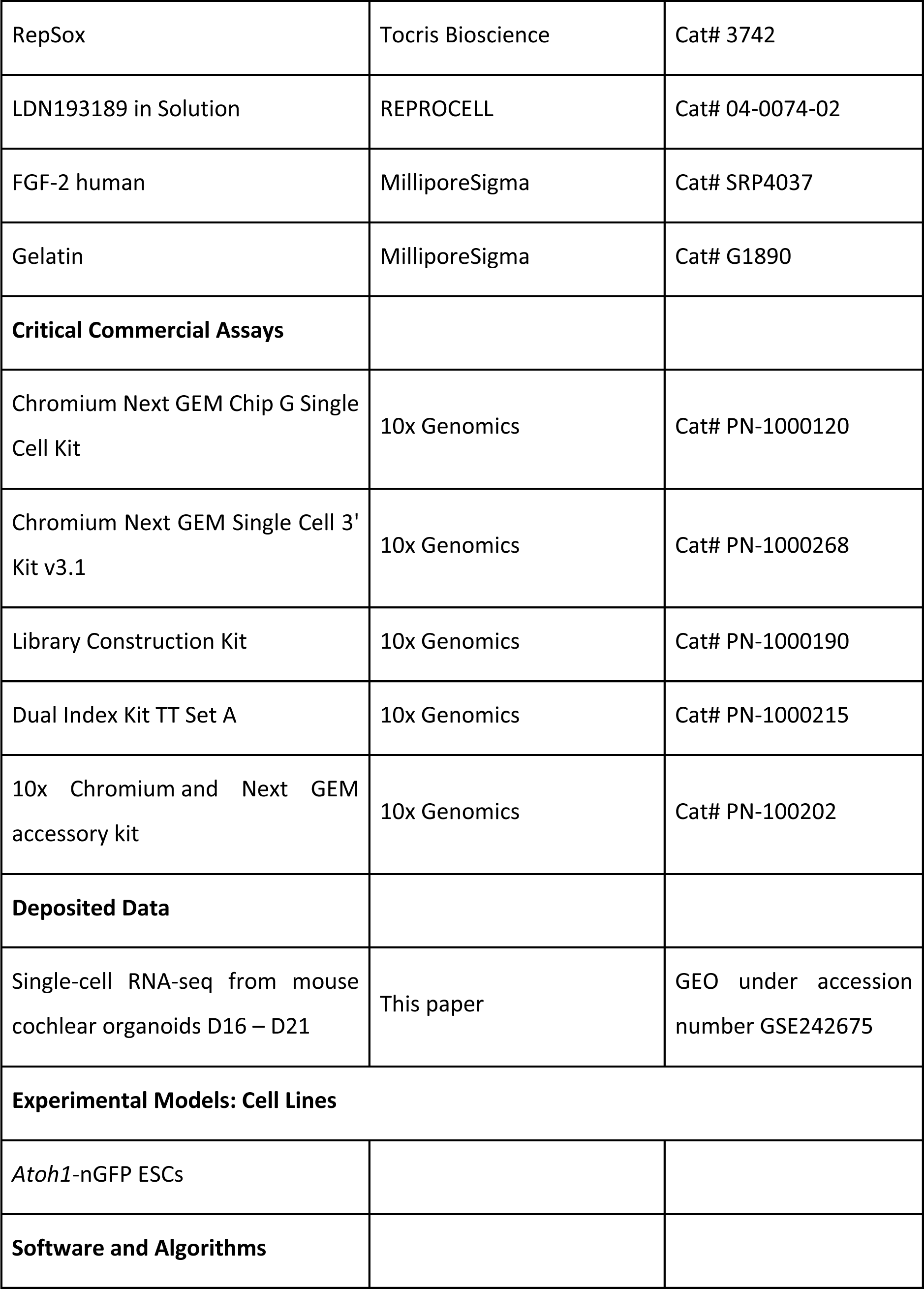
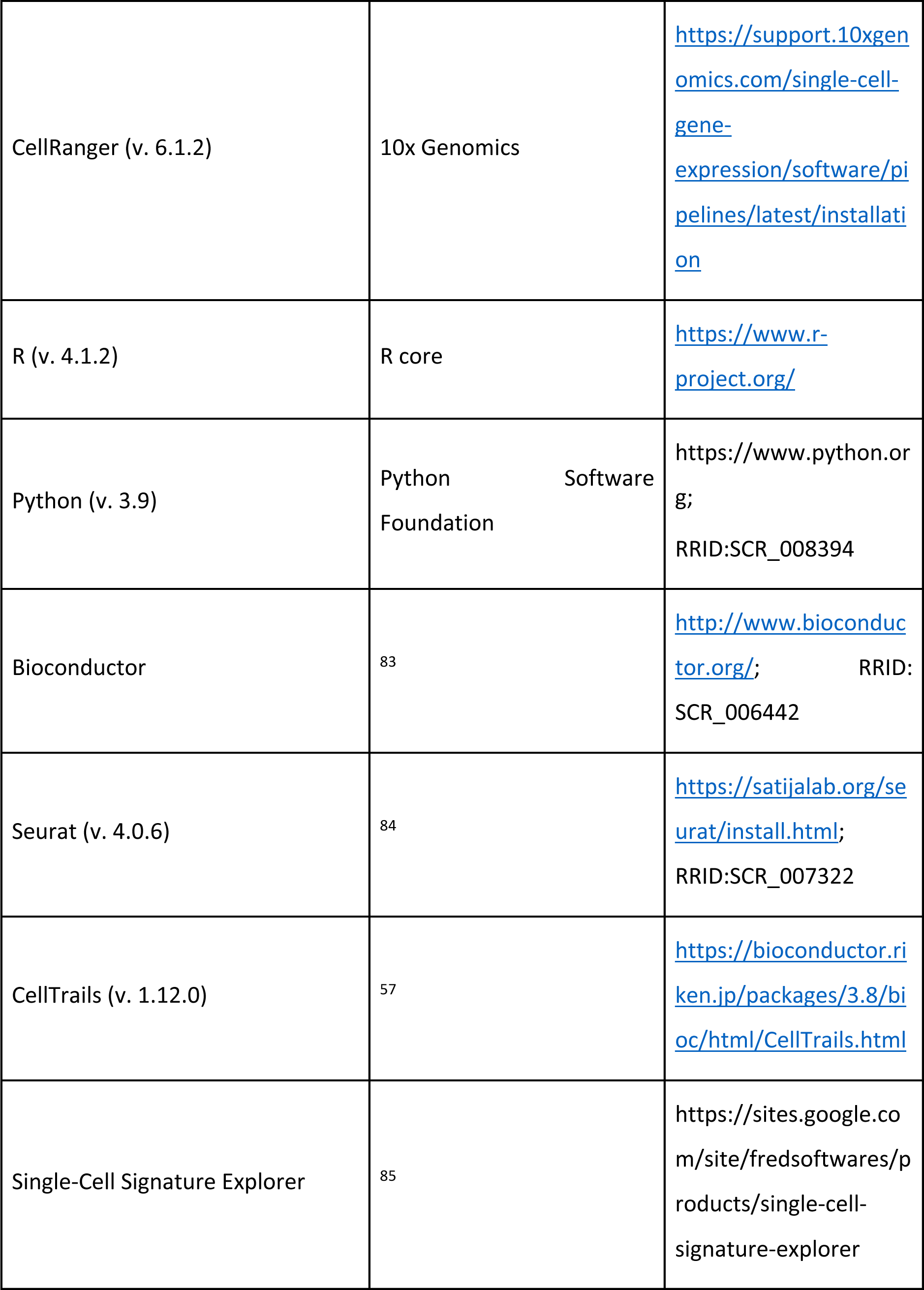
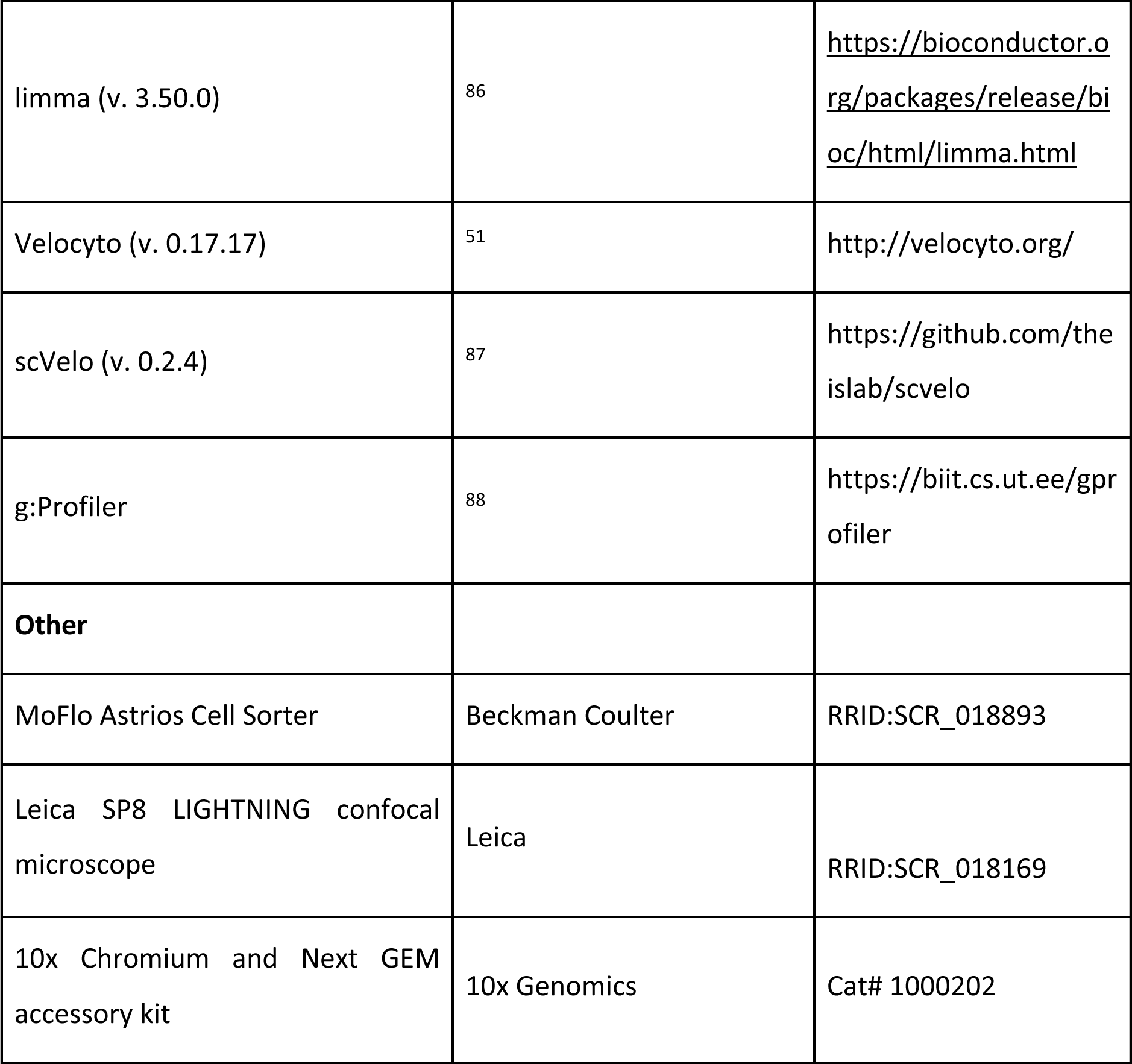

### Embryonic stem cell culture

Embryonic stem cells (ESCs) from transgenic Atoh1/nGFP reporter mice^25^ were obtained as a gift from Dr. Stefan Heller, Stanford University.^89^ The ESCs were maintained under serum-free, feeder-free conditions in a humidified incubator with 5% CO2 at 37°C. The cells were cultured in 2i-LIF media composed of a 1:1 mixture of Advanced DMEM/F12 and Neurobasal supplemented with 1X B27 without vitamin A, 0.5X N2, 1X GlutaMAX, 1000 U/mL LIF, 3 μM CHIR99021, and 1 μM PD0325901. Cells were passaged every 2-4 days using TrypLE, returning to frozen stocks after reaching a maximum of 25 passages.

### Inner ear organoid production

The inner ear organoids were generated essentially as described previously.^90^ On day 0 (D0), the ESCs were dissociated into a single-cell suspension using TrypLE (Thermo Fisher) and aggregated into spheroids by placing 3,000 cells per well round-bottomed 96-well Nunclon Sphera Microplates containing GMEM, 1.5% KnockOut Serum Replacement, 15 mM HEPES, 1X nonessential amino acids, 1 mM sodium pyruvate, and 0.1 mM β-mercaptoethanol. On D1, the media was supplemented with a final concentration of 2% v/v Growth Factor Reduced Matrigel (referred to throughout as Matrigel). On D3, 10 ng/mL BMP-4 and 1 μM RepSox were added to each well followed by the addition of 1 μM LDN193189 and 100 ng/mL FGF-2 on D4.25. On D8, aggregates were transferred to a new 96-well microplate with maturation media (MM) containing advanced DMEM/F12, 1x N2, 15 mM HEPES, and 1x Glutamax with 1% v/v Matrigel and 3 μM CHIR99021. Starting from D10, half of the media was exchanged with fresh MM every day until the specimens were required for downstream applications.

### Single cell isolation and flow sorting

Whole inner ear organoids from day 16 to 21 of the *Atoh1*-nGFP ESC line were processed for single cell preparation as previously described ^91^. To enrich for inner ear HCLCs, we purified cells with FACS with a MoFlo Astrios instrument (Beckman Coulter, University of Michigan Flow Cytometry Core). These samples were then used for standard 10x Genomics preparations for scRNA-seq experiments.

### 10x Genomics protocol

Single-cell processing and next-generation sequencing were performed at the Advanced Genomics Core at the University of Michigan. Sequencing was performed with a 10x Chromium and Next GEM accessory kit (10x Genomics, 1000202) and Chromium Next GEM Chip G Single Cell Kit (10x Genomics, 1000120) for scRNA-seq. The following kits were used for library preparation: Chromium Next GEM Single Cell 3’ Kit v3.1 (10x Genomics, 1000268), Library Construction Kit (10x Genomics, 1000190), and Dual Index Kit TT Set A (10x Genomics, 1000215).

### scRNA-seq analysis

The scRNA-seq datasets was analyzed using the Seurat v3 pipeline.^26^ First, we selected cells with the number of features ranging from 600 to 8000, and the maximum allowed fraction of mitochondrial genes per cell was set to 10%. A total of 15001 cells passed the quality control for further analysis. After the pre-processing step, log normalization was performed, and the top 3,500 highly variable genes were identified with the *vst* method with default settings. To avoid the domination of highly expressed genes, we scaled the datasets and applied PCA to reduce dimensionality. The first 15 principal components were chosen to construct the shared nearest neighbor graph with 25 nearest neighbors (*k.param=25*). After the optimization of the shared nearest neighbor modularity, clusters of cells were identified using *FindClusters* function with resolution 0.6. We leveraged UMAP to visualize the scRNA-seq clustering results and metadata information, such as sample ID. To determine cell identities for each cluster, we first identified DEGs for each cluster with the *FindAllMarkers* function in the Seurat package with the following parameters: *only.pos=TRUE*, *min.pct=0.25*, *logfc.threshold=0.25*, *test.use=“wilcox”*. Adjusted Bonferroni-corrected P-values of 0.05 were used for multiple testing correction. We determined cell identities by comparing cluster specific DEGs with published canonical marker genes.

### RNA Velocity Analysis

To calculate the RNA velocity of single cells, we used CellRanger output BAM file and executed the Velocyto run10x command. This generated a loom file containing the spliced and unspliced reads for each cell. Each dataset’s reads were separately quantified and subsequently merged after quantification. The resulting merged loom file was read into python. We collected cluster annotations from the Seurat object and retained only HCLCs and SCLCs. Next, we preprocessed the subset clusters using scVelo to perform filtering, normalization, and nearest neighbor assignment. The unspliced and spliced reads were averaged within the neighborhood by scVelo’s pp.moments method with 30 principal components among 50 neighbors. Velocities were obtained using scvelo.tl.velocity with default parameters, followed by projecting the velocity inference onto UMAP using pl.velocity embedding grid.

### Enrichment Score Analysis

We employed Single-Cell Signature Explorer to calculate enrichment scores. Single-Cell Signature Explorer computes enrichment scores of pre-defined gene set across single cells and enables visualization of scores on a UMAP plot. We determined differentially expressed genes between two groups samples and used them as gene set to calculate the signature score using normalized UMI counts of genes.

### Auditory vs vestibular cell types

First, we identified DE genes between auditory and vestibular HCs from a microarray dataset^45^ using limma (v3.50.0) package. Probes not detected in the data were filtered out. The filtering criteria required detection P value < 0.01 in at least two samples and this left 23051 probes. Arrays were then normalized using quantile normalization. Linear modelling was performed with contrasts fit to identify differences between auditory and vestibular HCs. An empirical Bayes moderation of the standard error was applied, and t-tests were used to assess significance accompanied by Benjamini–Hochberg correction for multiple testing. DEGs were filtered by adjusted P value < 0.05. To find unique gene sets expressed exclusively by auditory or vestibular HCs, we only kept DEGs that present in HCs but not in SCs. The same analysis was performed with auditory and vestibular SCs. In the end, we detected 354 auditory HC specific genes, 307 vestibular HC specific genes, 397 auditory SC specific genes and 449 vestibular SC specific genes which were used as gene sets in Single-Cell Signature Explorer.

### Inner vs outer hair cells

Lists of genes enriched in either IHCs or OHCs of neonates and adults were obtained from previously published data.^66^ There are 806 IHC-enriched genes of P0, 1045 IHC-enriched genes of adult, 659 OHC-enriched genes of P0 and 175 OHC-enriched genes of adult, respectively. Each list was used as input to calculate the signature score in each single cell and visualized in the UMAP. To enable comparisons between groups, we normalized scores by dividing the size of each gene set.

### Type I vs II vestibular hair cells

An *in vivo* generated HC dataset^7^ was used to compute DEGs of extrastriolar Type I, Type II and striolar HCs. We employed Seurat’s *FindAllMarkers* function to detect DEGs of each group with the following parameters: *only.pos=TRUE*, *min.pct=0.25*, *logfc.threshold=0.25*, *test.use=“wilcox”*. P value was adjusted for multiple testing with Bonferroni correction. Genes with adjusted p-values < 0.05 were kept in gene set. The Number of genes of extrastriolar Type I, II HCs and striolar HCs are 1207, 1233 and 548, respectively. Enrichment scores were visualized in both UMAP and trajectory graph.

### Joint Analysis

To conduct joint analysis of *in vitro* generated HCLCs, neonatal utricle and P2 organ of Corti, we followed Seurat v3 standard data integration pipeline. This approach applies canonical correlation analysis (CCA) to identify anchor cells between pairs of datasets. Briefly, 3500 highly variable genes were used for finding alignment anchors and top 25 dimensions from CCA were used for defining neighbor search space, with the function *FindIntegrationAnchors*. These anchors, determined and scored for all sample pairs, were then used to integrate data across datasets using *IntegrateData* function. The Integrated data was scaled and reduced using PCA. We used UMAP for data visualization.

### Vestibular HC Trajectory Reconstruction

We applied CellTrails to reconstruct HC developmental trajectory using normalized scRNA-seq data. CellTrails employs a non-linear dimensionality reduction and hierarchical clustering method for identification of latent spaces to determine cell states and to infer pseudotime expression dynamics. To find a low-dimensional representation, we applied spectral embedding function *embedSample* with default parameters, resulting in 12 latent variables for further analysis. Then, hierarchical clustering with a *post-hoc* test was performed using the function *findStates* with following parameters: *min_size=0.02, min_feat=5, max_pval=1e-4, min_fc=1.8*. This identified 8 distinct cell states. Finally, we aligned individual cells onto the trajectory using *fitTrajectory* function.

### Similarity Matrix

To test two potential mechanisms conferring tonotopic identity, we calculated pairwise cell similarity scores between the *in vitro* and *in vivo* generated HCs. Specifically, we first generated metacells to denoise the dataset by aggregating the six nearest neighbor cells for both *in vitro* and *in vivo* data. By using the GEGs, we calculated the Euclidean distance for each pair of metacells between *in vitro* and *in vivo*. Next, several methods were used to convert the distance matrices into similarity matrices. For the Euclidean distance matrix, the traditional inverse method (1/(1+distance)) or radial basis function (exp⁡(-〖distance〗^2/(2σ^2))) was used to generate the similarity matrix. σ, bandwidth, was a hyperparameter. In our analysis, we defined σ^2=4. For cosine distance matrix, similarity scores were calculated as 1-distance. Finally, the similarity matrix was visualized with a heatmap. Metacells of both groups were ordered along developmental states derived by CellTrails.

### Cryosection and immunostaining

Aggregates progressing through the organoid differentiation protocol were collected at time points indicated and fixed in 4% paraformaldehyde (PFA) for 30-60 minutes at room temperature. The specimens were cryopreserved in increasing concentrations of sucrose (10%, 20%, and 30%) for 30 minutes each and stored overnight in a 1:1 mixture of 30% sucrose and Tissue-Tek O.C.T. (Thermo Fisher). Up to 25 aggregates per culture were transferred to cryomolds with O.C.T. and frozen on dry ice. Serial sections, 10 μm in thickness, were collected on SuperFrost-Plus slides (Thermo Fisher) and stored at −80°C until use. In some cases, single-cell suspensions of organoid cells were also prepared for immunostaining. Organoids were digested as above and added to SuperFrost-Plus slides containing phosphate buffered saline (PBS), either before or after flow cytometry. Cells were allowed to settle and adhere to the slide for 15 minutes before being fixed in 4% PFA for 30 minutes at room temperature.

Slides of cryosections or isolated cells were blocked in PBS with 10% normal donkey serum and 0.1% Triton X-100 and then incubated overnight at 4°C in primary antibodies diluted in PBS with 5% normal donkey serum and 0.05% Triton X-100. Primary antibodies included rabbit anti-Six1 (Sigma, 1:1000), rabbit anti-Pax2 (Invitrogen, 1:100), rabbit anti-MYO7A (Proteus, 1:200), mouse anti-SOX2 (Santa Cruz, 1:100), and rabbit anti-GFP (Millipore, 1:500). The samples were counterstained with AlexaFluor secondary antibodies, Alexa-conjugated phalloidin to label actin-rich hair bundles, and/or Hoechst 33242 to label the nuclei. The specimens were mounted in ProLong Gold Antifade solution and imaged under epifluorescence with an Olympus BX51 microscope and ORCA R2 CCD camera or using confocal microscopy with a Leica TCS SP5 system. Images were post-processed in MetaMorph (Molecular Devices) or ImageJ.^92^

### Statistics and reproducibility

Statistical analysis was performed using R on RStudio. A two-sided P value was considered statistically significant if *P <0.05, **P <0.01, and ***P <0.001.

## Data availability

All scRNA-seq raw and processed sequencing data generated in this study have been deposited to the NCBI Gene Expression Omnibus database and can be retrieved using the accession number GSE242675. Source data are provided with this paper.

## Code availability

The codes for key computational analyses are available on GitHub (https://github.com/waldhaus/InVitro_Hair_Cells). All of the packages used are available online.

## Acknowledgements

We thank the staff at the University of Michigan Advanced Genomics Core for help and support. This project was supported by funds from the Department of Veterans Affairs (I01 RX003400 to RKD).

## Author Contributions

JW, JL, and RKD conceived the study, and supervised the project. LJ and LL acquired and analyzed data. JW wrote the manuscript with the help of LJ, LL, JL, and RKD.

